# A Unified Mechanism for Depth Perception and Figure-Ground Organization

**DOI:** 10.64898/2025.12.10.693427

**Authors:** Tianlong Chen, Xuemei Cheng

**Author notes:** Corresponding author: Tianlong Chen.

## Abstract

Understanding how the visual system unifies depth perception and figure–ground organization remains a central challenge in vision science. Stereoscopic disparity and border ownership have long been treated as distinct processes, yet both arise with similar timing in area V2 and exhibit strikingly parallel properties. Here we propose a unified mechanism that integrates both within a single hierarchical framework.

Building upon the layered disparity representation introduced in our prior work, we show that **border ownership—serving as the key component of figure– ground organization—emerges intrinsically from thresholded relative-disparity differencing**, providing a unified basis for contour-based segmentation and depth perception. Using biologically constrained network models (TcNet), we demonstrate that this mechanism generalizes across binocular and illusory depth domains, bridging physical and non-epipolar disparity cues.

At the systems level, we identify complementary **dual feedback pathways from area V4**: V4→V1, which refines and re-weights layered disparity maps, and V4→V2, which integrates global contextual priors such as shape continuity and category consistency. Together, these recurrent loops balance fine-scale depth precision with large-scale figure–ground coherence, offering a unified explanation for perceptual alternations such as the Rubin Face–Vase and Kanizsa illusions.

**This framework integrates intrinsic disparity computation, border-ownership dynamics, and hierarchical feedback modulation into a coherent, biologically plausible model**. It provides a mechanistic account of how early visual areas interact with higher cortical feedback to achieve stable yet flexible perception of depth and figure–ground organization within a unified temporal cycle.

## 1. Introduction

A central challenge in visual neuroscience is understanding how the brain simultaneously encodes depth and figure–ground organization. Two phenomena lie at the heart of this challenge: *relative disparity (RD)*, which represents depth ordering between objects, and *border ownership (BO)*, which assigns a contour to one side of an object rather than the other.

Physiological studies indicate that both relative-disparity–based depth-order signals and border-ownership (side-of-figure) signals arise rapidly in V2. Border-ownership modulation appears approximately 25 ms after the onset of the classical V2 response (Zhou et al., 2000), and relative-disparity signals are expressed within the earliest phase of the V2 response as well, with surround-dependent modulation evident essentially from response onset (Thomas et al., 2002; Qiu & von der Heydt, 2005). These observations raise the long-standing question of *whether the two phenomena reflect deeply coupled or even unified computational mechanisms*.

In our previous work (Chen et al., 2025, *Understanding and Simulating Border Ownership Centered Segmentation*), we proposed a model of figure-ground in which BO is shaped by global context awareness and layered disparity channels. Though that study treated disparity and BO as related but distinct cues, our further analysis suggests a more fundamental relationship: ***BO may not merely co-occur with RD, but may actually arise as a thresholded form of RD itself***.

This perspective shifts the theoretical foundation of BO, framing it not as a late or context-dependent assignment but as an *intrinsic transformation of disparity*, computed locally and expressed with short latency. Such a view provides a parsimonious explanation for decades of neurophysiological findings across classical, naturalistic, and illusory stimuli, while offering ***a unified account in which depth perception and figure–ground organization emerge as two expressions of the same underlying computation***.

### Paper Organization

**Section 2** reviews background on disparity, border ownership, and their long-standing separate treatment in vision science.

**Section 3** revisits prior work on relative and border ownership, including our earlier models, and motivates the hypothesis of a deep computational equivalence between them.

**Section 4** formalizes this hypothesis by introducing local differencing kernels and constructing relative-disparity and border-ownership representations.

**Section 5** presents empirical comparisons between opposite-channel (OC) and relative-disparity (RD) coding across disparity-defined, contrast-defined, and illusory datasets.

**Section 6** integrates these results into a unified theoretical account, examining the nature of epipolar and non-epipolar disparity, layered disparity representation, global-context modulation, and category selectivity, and the roles of V4 feedback.

**Section 7** discusses limitations, open questions and future directions.

**Section 8** concludes with a summary of the unified depth perception and figure-ground mechanism proposed in this work.

## 2. Background

### 2.1. Disparity and Depth Perception

Binocular disparity is the primary cue by which the visual system recovers 3-D structure.

*Absolute disparity*, computed in V1, encodes the offset of retinal images relative to fixation. Although fundamental, absolute disparity is reference-dependent and limited in its perceptual range (Cumming & Parker, 1999).

By contrast, *relative disparity (RD)*, computed in V2, represents the depth difference between surfaces and provides a stable, fixation-invariant basis for depth ordering (Thomas, Cumming & Parker, 2002). RD is widely recognized as the dominant neural signal supporting judgments of depth arrangement in both natural and controlled settings (Parker, 2007). Because RD directly represents depth relations rather than raw binocular geometry, it plays a crucial role in perceptual stability across gaze shifts and viewing conditions.

### 2.2. Border Ownership

Border ownership (BO) signals, first characterized in macaque V2, specify which side of a contour belongs to an object. Seminal studies demonstrated neurons whose responses depend on the side of figure relative to a border (Zhou et al., 2000). Subsequent work showed that these signals are strongly influenced by global configuration, long-range relationships, and surface context, indicating integration well beyond the classical receptive field (Qiu & von der Heydt, 2005).

BO modulation arises rapidly—approximately 25 ms after the onset of the classical V2 response—indicating that ownership signals are initiated very early in V2 processing (Zhou et al., 2000; Craft et al., 2007). Contextual grouping and surface-continuity cues further shape and stabilize BO signals over subsequent processing stages (Craft et al., 2007; Williford & von der Heydt, 2013; Williford & von der Heydt, 2016), helping maintain coherent figure–ground interpretation under ambiguous or dynamic conditions.

### 2.3. The Unresolved Relationship

Despite their similar timing and shared locus in V2 (Thomas et al., 2002; Zhou et al., 2000), relative disparity (RD) and border ownership (BO) have historically been treated as distinct processes. RD has typically been framed as a depth-ordering signal derived from binocular geometry (Poggio, 1995; Roe et al., 2007), whereas BO has been framed as a contour-ownership signal shaped by global scene context and long-range integration (Von der Heydt, 2015; Chen et al., 2025).

Yet several observations indicate that the two phenomena may be more deeply linked than traditionally assumed. BO can be generated even in the absence of explicit disparity, as in illusory contours and Kanizsa figures (Kanizsa 1976; Von der Heydt et al., 1984; Chen et al., 2025). Conversely, Qiu et al., (2005) demonstrated that, for many V2 neurons, the *preferred side of figure coincided with the near side of the neuron’s preferred stereoscopic depth*. This alignment provides *direct physiological evidence* that depth-order signals and contour-ownership preferences *co-exist within the same V2 population*, suggesting that they may arise from a shared underlying computation.

Together, these findings motivate a re-examination of whether RD and BO might be different expressions of a single underlying computation, rather than independent cues combined at a later processing stage.

## 3. Motivation and Neurophysiological Basis

### 3.1. Neurophysiological Studies

Neurophysiological research has identified distinct but closely related signatures of disparity and border ownership in early visual cortex.

Cumming & Parker (1999) showed that *absolute-disparity* signals are strongly represented in V1, providing some of the earliest binocular depth information.

Building on this, Thomas et al. (2002) demonstrated that many V2 neurons express *relative-disparity* sensitivity from the early phase of their responses, reflecting surround-dependent modulation beyond absolute disparity tuning.

Complementary work by Qiu and von der Heydt (2005) and Williford and von der Heydt (2013) revealed robust *border-ownership (BO)* signals in V2: neurons code the side of figure even when the local edge stimulus is unchanged, and this selectivity depends on both local edge orientation and global scene context.

Crucially, Zhou et al. (2000) reported that border-ownership modulation in V2 appears with very short delay—on the order of *∼25 ms after response onset*— indicating that BO emerges early in V2 processing.

Taken together, these findings show that relative disparity and border ownership are both computed in V2 and manifest in closely overlapping timing windows. The observed correspondence between preferred depth sign and preferred side-of-figure in many V2 neurons (Qiu & von der Heydt, 2005) further *suggests that the two signals may be more tightly linked than traditionally assumed*.

### 3.2. Modeling Approaches

Computational models of disparity and border ownership have advanced substantially over the past decades, yet almost always as *separate theoretical lines*. Here we review the major model families relevant to each domain and highlight the conceptual limitations that motivate a unified mechanism.

#### Disparity Models

The *binocular energy model* has been the dominant framework for explaining disparity selectivity. The seminal work of Ohzawa, DeAngelis, and Freeman (1990, 1997) demonstrated that disparity tuning can arise from *phase* differences between left- and right-eye receptive fields.

Anzai, Ohzawa, and Freeman (1999) expanded this framework by showing that both *phase disparities* and *position disparities* contribute to disparity selectivity in V1, establishing the canonical basis for *absolute disparity* processing.

To account for *relative disparity*, later extensions of the energy model incorporated multi-region pooling, center– surround mechanisms, or paired receptive fields (Cumming et al., 1999; Thomas et al., 2002). Roe et al., (2007) reviewed how such mechanisms could support first- and second-order disparity selectivity, explaining percepts such as surface slant and curvature.

While these approaches capture binocular geometry, they do not directly address contour assignment, ownership direction, or figure–ground structure.

Beyond the energy framework, Grossberg & Yazdanbakhsh (2005) proposed a laminar boundary– surface architecture in which left- and right-eye monocular boundaries are computed independently and subsequently fused to form binocular boundary representations supporting 3D surface perception, stratification, and neon color spreading.

Although this model links disparity with surface formation, the fused boundaries are unsigned and do not encode border-ownership polarity, nor does the model propose a local computation that unifies relative disparity with contour assignment.

Relative disparity is classically defined as the difference between the absolute disparities of two points (Howard & Rogers, 1995). Recent psychophysical work (Ding et al., 2024) provides the latest compelling support for this formulation, demonstrating that the relative disparity between two points P_i_ and P_j_ is given by *d*_*ij*_ = *d*_*i0*_ – *d*_*j0*_, where each absolute disparity d_i0_, d_j0_ is measured with respect to fixation. This formulation highlights the noise-canceling property of relative disparity: global offsets cancel, enhancing sensitivity to local depth differences.

These findings reinforce the view that simple directed differencing may lie at the core of relative-disparity computation in the visual system.

#### Border Ownership Models

In contrast, models of border ownership have been developed largely independently of disparity, focusing on contour assignment and contextual interpretation.

The influential *grouping-cell model* by Craft et al. (2007) posits specialized grouping cells (g-cells) that integrate contextual signals from border-ownership– selective edge units (b-cells) and bias ownership assignment toward one side of a contour. In this framework, border ownership arises from a combination of *horizontal long-range interactions and top-down feedback from higher areas such as V4*, which help enforce global configuration and shape cues.

However, disparity is not represented in this architecture, and the model does not incorporate depth cues.

A contextual-interaction account was proposed by Zhaoping (2005), who argued that border ownership could be generated *within V2* through intracortical interactions among orientation-selective neurons, without requiring higher-level feedback. In this model, local recurrent dynamics and lateral competition bias one side of an edge toward figure assignment.

Subsequent discussions by Zhaoping (2011) further emphasized that V1 provides strong contextual modulations and may contribute proto-figure signals, but did not propose that V1 itself computes border ownership. The notion that V1 might generate BO remains controversial, as most physiological evidence indicates V2 as the primary locus of robust, polarity-specific border-ownership selectivity.

The *incremental grouping model* by Roelfsema et al. (2006, 2011) frames border ownership as part of a larger process of perceptual grouping, in which contour fragments are bound together iteratively through attentional feedback and recurrent horizontal integration. While powerful for explaining scene segmentation, this framework treats BO as an independent contour-binding process, rather than a direct consequence of disparity computation.

Finally, Grossberg (2016) extended the *laminar boundary–surface framework* to incorporate border ownership, stereoscopic cues, and Gestalt grouping rules within a unified laminar architecture.

Although this model integrates BO with disparity and context, it retains distinct mechanisms for boundary grouping and BO assignment and does not propose a single feedforward operation that simultaneously yields both relative disparity and ownership polarity.

#### Synthesis

Overall, disparity models focus on binocular geometry and depth ordering, whereas border-ownership models emphasize contour assignment, contextual grouping, and scene interpretation.

*No existing framework directly unifies relative disparity and border ownership through a single local computation*.

### 3.3. Our Prior Model

In our earlier work (*Border Ownership, Category Selectivity, and Beyond*, Chen et al., 2022), we introduced a biologically inspired representational scheme for border-ownership and category-selectivity, together with TcNet, a neural network designed to jointly generate border ownership and category selectivity without relying on T-junction cues or top-down guidance. This framework served as a foundation for modeling figure– ground organization and was validated on monocular stimuli.

In our more recent work (*Understanding and Simulating Border Ownership-Centered Segmentation*, Chen et al., 2025), we extended this framework to broader vision scenarios, demonstrating consistent performance across contrast-defined, disparity-defined, illusory, and contour-defined objects. These results supported the idea that a common coding strategy may underlie figure– ground assignment across diverse stimulus classes.

Building on TcNet, we further proposed an integrated segmentation framework—referred to as the *Border Ownership–Centered Figure–Ground Model* (Chen et al., 2025)—which combined *layered disparity channels* (epipolar/near, logarithmic/far, and additional channels such as optic flow) with an *active-neuron mechanisms* for surface filling-in. This architecture provided a biologically motivated way to link absolute disparity and border ownership within a unified segmentation pipeline.

However, in all prior formulations, disparity and border ownership were still treated as *cooperating but distinct* computational modules.

### 3.4. Revisiting Our Prior Findings

A key experiment in our prior work (Chen et al., 2025, Section “*Illusory Objects and Illusory Contours*”), involved applying monocular TcNet01 to illusory Kanizsa figures. We observed that adding synthetic absolute (Near) disparity degraded border-ownership performance, leading us at the time to conclude that explicit depth cues were *not required* for border-ownership generation in monocular or binocular illusions.

A closer re-examination of that result, however, points to a different and more consequential interpretation—one that ultimately motivated the present framework. Rather than indicating independence from depth, the finding suggests that *illusory border-ownership signals themselves may encode implicit depth relations*. That is, even without physical (absolute) disparity, the visual system may infer *relative-disparity-like structure* from the illusory configuration, generating depth-order information that is functionally similar to stereoscopic relative disparity.

This reinterpretation is consistent with Coren‘s (1972) proposal that *both monocular and binocular subjective contours derive from latent or inferred depth cues*. Under this view, the Kanizsa border ownership signal is *not depth-free*; instead, it *embeds a depth-order inference* that behaves analogously to relative disparity.

If this perspective is correct, then our earlier findings suggest a deeper conclusion and the ***central hypothesis*** of this work: ***border ownership and relative disparity may not merely interact—they may be computationally equivalent expressions of the same underlying operation***.

## 4. Relative Disparity and Border Ownership Representation

### 4.1. Rationale for Relative Disparity

Neuroscientific and psychophysical evidence suggests that relative disparity is computed as the difference between absolute disparities of two spatially distinct points. Thomas et al. (2002) demonstrated that V2 neurons are sensitive to depth differences between “regions”, while Roe et al. (2007) reviewed how specific receptive-field arrangements could support such differencing operations. More recently, Ding et al., (2024) provided compelling psychophysical support that relative disparity can be formally expressed as:

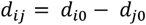

where *d*_*i0*_ and *d*_*j0*_ denote the absolute disparities of two points measured with respect to fixation.

However, two fundamental questions remain unresolved.

First, *when* in the visual processing hierarchy relative disparity is explicitly computed, particularly in relation to the emergence of border ownership, remains unresolved. Absolute disparity signals emerge in V1 (Cumming et al., 1999), whereas relative disparity selectivity is present from the earliest phase of V2 responses (Thomas et al., 2002).

Second, *how* the visual system selects the specific “two points” whose disparities should be differenced is unclear. The operation might arise from fixed receptive-field structure, spatial pooling across local neighborhoods, or dynamic context-dependent selection.

If the hypothesis of a deep computational equivalence between relative disparity and border ownership is correct, then relative disparity cannot be meaningfully defined over pre-segmented objects or regions, because object structure itself depends on border ownership. Instead, relative disparity must be formed *concurrently with* the emergence of border ownership.

Under this view, the most biologically plausible form of relative disparity is the differencing of disparities between neighboring units (pixels or neurons) ***uniformly across*** the visual field, rather than between preestablished surfaces or regions.

### 4.2. Kernel-based Implementation

Based on the hypothesis that relative disparity emerges from ***local, uniform spatial differencing*** of absolute disparities, we propose two possible kernel sets (denoted *k3*.*4* and *k1*.*5*) for transforming absolute disparity maps into relative disparity maps.

#### The k3.4 Kernel Set

The *k3*.*4 kernels* are directly inspired by the border-ownership ground truth generation used in Chen et al. (2022) and Chen et al. (2025), where ownership is determined by the depth difference across a border point. The same logic can be applied to approximate local relative disparity. The *k3*.*4 kernel set* consists of a row kernel *K*_*r*_ and a column kernel *K*_*c*_ (illustrated in Figure 1):

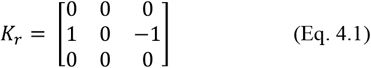

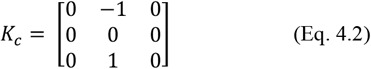

**Figure 1.**
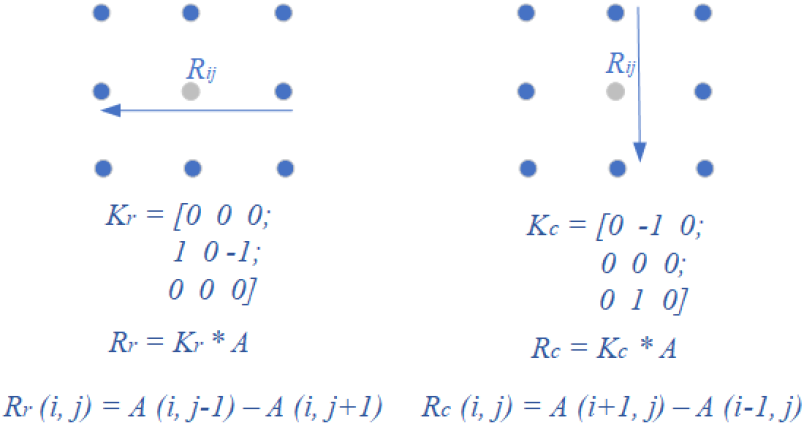
Visualization of the k3.4 Kernel Set. The relative disparity ***R***(***i, j***)at pixel location (***i, j***)is computed by contrasting disparities on opposite sides along the horizontal or vertical axis, capturing local disparity contrast in a symmetric fashion.

#### The k1.5 Kernel Set

The *k1*.*5 kernels* are motivated directly by the formal definition of relative disparity itself, as shown by Ding et al., (2024), in which relative disparity is computed as the difference between two sampled disparities. This leads to compact one-sided kernels (illustrated in Figure 2):

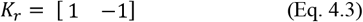

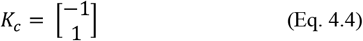

**Figure 2.**
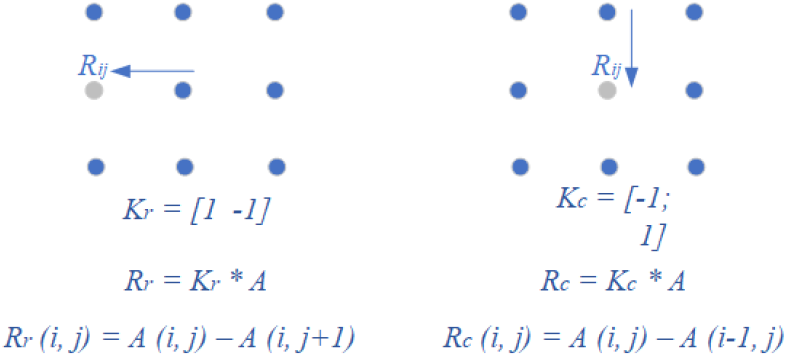
Visualization of the k1.5 Kernel Set. The relative disparity ***R***(***i, j***)at pixel location (***i, j***)is computed by differencing each pixel’s disparity with its immediate rightward (row) or upward (column) neighbor. This implements the minimal, locally defined differencing scheme.

#### Relative Disparity Maps

Applying convolution (∗)with these kernels produces horizontal and vertical relative disparity maps:

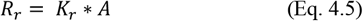

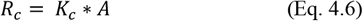

Here, *A* denotes the layered disparity representation introduced in Chen et al. (2025), which includes absolute disparities, optic-flow channels and other disparity-related components. *A* is represented as a tensor of size *M×N×C*, where *M×N* is the spatial resolution and *C* is the number of channels. *R*_*r*_ and *R*_*c*_ are the resulting relative disparity maps computed along the row and column directions, respectively.

#### Border Ownership Maps

In our prior work (Chen et al., 2022; Chen et al., 2025), border ownership was generated using TcNet, an encoder-decoder neural network trained to produce ownership maps, with borders subsequently extracted through thresholding of its raw outputs.

Inspired by that procedure, we follow an analogous strategy here: applying thresholding to the relative disparity maps *R*_*r*_ and *R*_*c*_ obtained above to derive border-ownership maps.

For each disparity channel *c* ∈ {1, …, *C*} in the layered disparity representation *A*, we compute channel-specific relative disparity maps along both the row and column directions. Thresholding is then applied to these channel-wise maps to identify candidate ownership contours.

The final border-ownership contour maps are then obtained by *summing thresholded relative disparity maps across all disparity channels*, thereby producing a two-component vector field encoding owner-side direction, as follows:

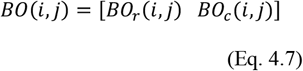

With

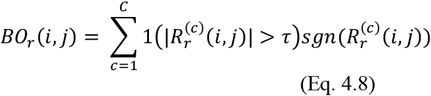

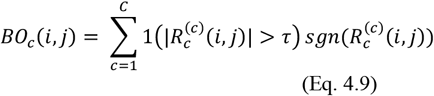

where:

- 1(⋅)is the indicator function with threshold τ, returning 1 when the condition holds and 0 otherwise;
- *sgn*(⋅)denotes the signum function, returning 1 if positive, −1 if negative, and 0 otherwise;
- 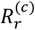and 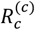 denote the relative-disparity maps computed along the row and column directions for channel *c*; and
- *BO*(*i, j*)is the resulting border-ownership at pixel (*i, j*)whose row and column components *BO*_*r*_ (*i, j*)and *BO*_*c*_ (*i, j*)follow the RD/BO configuration defined in Figure 4.

#### Relative Disparity and Border Ownership Representation

In Chen et al. (2022), we introduced the *opposite channel (OC) rule* for border-ownership coding, in which *occluding contour segments with opposite owner sides are assigned to separate channels*. This scheme provides an effective means of disambiguating figure–ground relationships.

Here, we extend this channel-based representation to relative disparity.

Specifically, relative disparity and its induced border ownership are separated into distinct channels *according to row vs column orientations, and are further subdivided by the sign of relative disparity*. We refer to this representation as *relative-disparity-based coding* (RD coding) (Figure 4).

As illustrated in Figure 3, the owner-side direction is determined jointly by *row/column orientation* and the *sign* of relative disparity computed from (Eq. 4.7).

**Figure 3.**
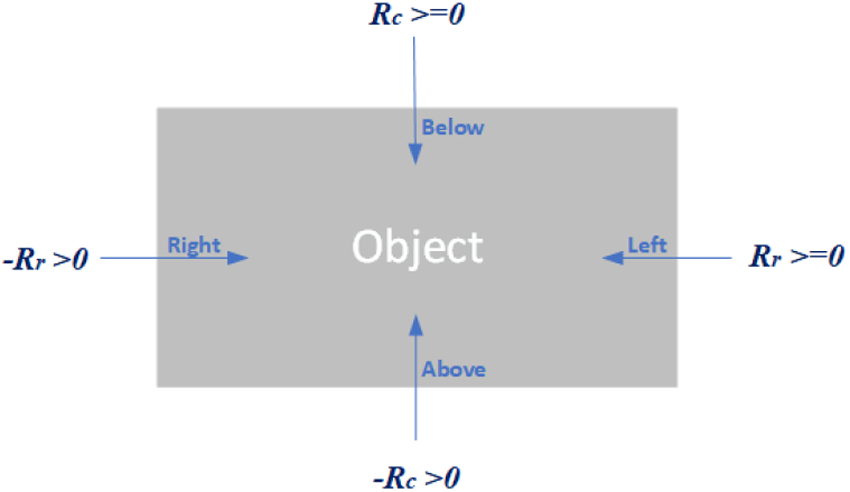
Visualization of Border Owner-side directions on a simple rectangle object according to row and column direction and sign of relative disparity. Arrows across borders labeled ‘below’, ‘left’, ‘above’, and ‘right’ point towards the border owner. Importantly, *the direction of increasing disparity is assigned as the owner-side direction, corresponding to the perceptually nearer region*.

**Figure 4.**
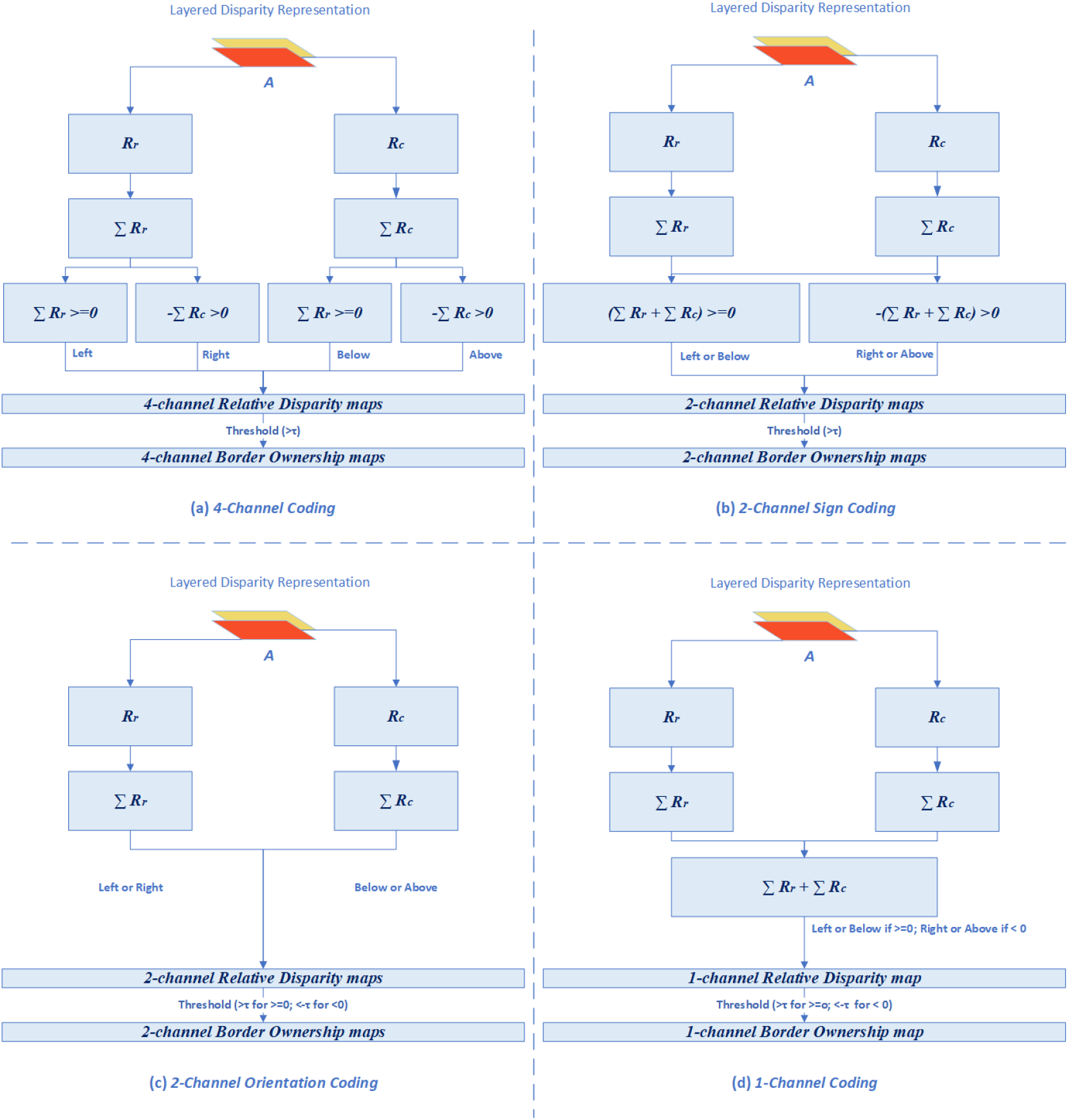
Relative-Disparity-based Coding for Relative Disparity and Border Ownership. Owner-side directions are indicated as ‘below’, ‘left’, ‘above’ and ‘right’. **(a) 4-channel coding**: four distinct channels represent relative disparity and border ownership by orientation and sign: row-positive (‘left’), row-negative (‘right’), column-positive (‘below’), and column-negative (‘above’). **(b) 2-channel sign coding**: two channels respectively encode the summed contributions of positive and negative disparities across both orientations. **(c) 2-channel orientation coding**: two channels separate relative disparity by orientation only (row and column). **(d) 1-channel coding**: a single channel combines summed row and column information with sign into a unified map. **Note**: These channels represent border-ownership-oriented outputs. In all configurations, summation may be performed *after* thresholding, which preserves intermediate RD maps for depth perception or downstream processes (e.g., dynamic surface filling-in) and is biologically more plausible. When summation occurs after thresholding, each BO-oriented channel corresponds to *C* underlying RD layers, where *C* is the number of disparity layers in the Layered Disparity Representation.

***Most importantly, the direction of the increasing disparity is assigned as the owner-side—****that is, the side perceptually nearer to the observer*.

Equivalently, *the sign of relative disparity* yields *the two opposite channels, directly paralleling the OC representation* used in our prior work (Chen et al., 2022; Chen et al., 2025).

To formalize the coding scheme, Figure 4 summarizes four possible relative-disparity/ownership channel configurations, mirroring the 2-channel and 4-channel BO representations introduced previously (Fig. 1 in Chen et al., 2022 or Fig. 2 in Chen et al., 2025). In all configurations, ***border ownership is obtained directly by thresholding relative disparity maps***, without requiring explicit contour templates, object models, or higher-level inference.

#### Testing the Central Hypothesis

To evaluate *the central hypothesis of this work*—that relative disparity and border ownership may share a deep computational equivalence—we compare the performance of the proposed relative-disparity-based (RD) coding with the established opposite-channel-rule (OC) coding (Chen et al., 2022; Chen et al., 2025). Although the two schemes arise from different conceptual motivations, their representations were deliberately aligned so that RD coding mirrors the channel organization of OC coding. This alignment enables a direct, controlled comparison on identical datasets.

Within this framework, OC coding serves as a rule-based benchmark for border-ownership assignment, whereas RD coding provides a disparity-driven substrate that could, in principle, give rise to the same owner-side assignments. Assessing how closely RD coding approximates OC coding across diverse conditions offers a direct test of the hypothesis that the two may reflect the same underlying computation.

In the next section, we move from theoretical formulation to empirical validation, presenting experiments that compare the two schemes across diverse datasets and conditions.

## 5. Experiments and Results

### 5.1. Brief Comparison of Coding Schemes

Opposite-channel-rule (OC) coding and relative-disparity-based (RD) coding represent distinct but partially aligned strategies for border ownership.

OC coding enforces exclusivity by assigning each border pixel to a single channel according to its owner side. In contrast, RD coding derives ownership from local disparity differencing, allowing pixels to contribute across both row- and column-based channels, with sign of relative disparity serving as the analogue of opposite-channel coding.

Despite this conceptual difference, RD coding was deliberately designed to mirror the channel organization of OC coding, enabling direct, controlled comparisons on identical datasets.

In this framework, OC coding provides a rule-based representation of figure–ground assignment, while RD coding offers a disparity-driven substrate that could, in principle, underlie such assignments.

As summarized in Figure 4, the four RD/BO configurations differ in how they treat orientation and sign:

- 4-channel configuration (Figure 4(a)) and the 2-channel orientation configuration (Figure 4(c)) keep row and column contributions strictly separated, with no overlap across orientations.
- The remaining two configurations permit row–column overlap, which can introduce partial cancellation depending on disparity structure.
- Among the four, the 2-channel sign configuration (Figure 4(b)) is the closest RD analogue to the default 2-channel OC coding widely used in our prior work (Chen et al., 2022; Chen et al., 2025).
- The 1-channel RD configuration (Figure 4(d)) functions as a secondary approximation to 2-channel RD sign coding (and therefore an approximation to 2-channel OC coding) when positive and negative values are separated into different output channels.

Accordingly, in the experiments that follow, we focus on the two most interpretable comparisons:

- 2-channel sign RD coding (Figure 4(b)), and
- 1-channel RD coding (Figure 4(d))

This pairing allows us to test whether the sign of relative disparity alone is sufficient to support opposite-owner-side assignment—one of the key mechanisms hypothesized to couple relative disparity and border ownership—and to evaluate whether RD coding can approximate OC coding across diverse datasets.

### 5.2. Datasets

For direct comparison between OC and RD coding, we used the same datasets as in our prior study (Chen et al., 2025). These include:

- **Random Dot Stereograms (RDS)** — disparity-defined objects (dataset publicly available at Chen, 2025a).
- **Modified Virtual KITTI 2 (VKitti)** — photorealistic contrast-defined scenes (dataset publicly available at Chen, 2025b; original VKitti from Cabon et al., 2020).
- **Kanizsa figures with synthetic Near disparity (Kanizsa)** — illusory-contour stimuli augmented with absolute Near disparity, representing the critical case that motivated an alternative interpretation of border ownership in our prior work (dataset publicly available at Chen & Cheng, 2026).

Representative samples of the three datasets used in the experiments are shown in Figure 5–Figure 7.

**Figure 5.**
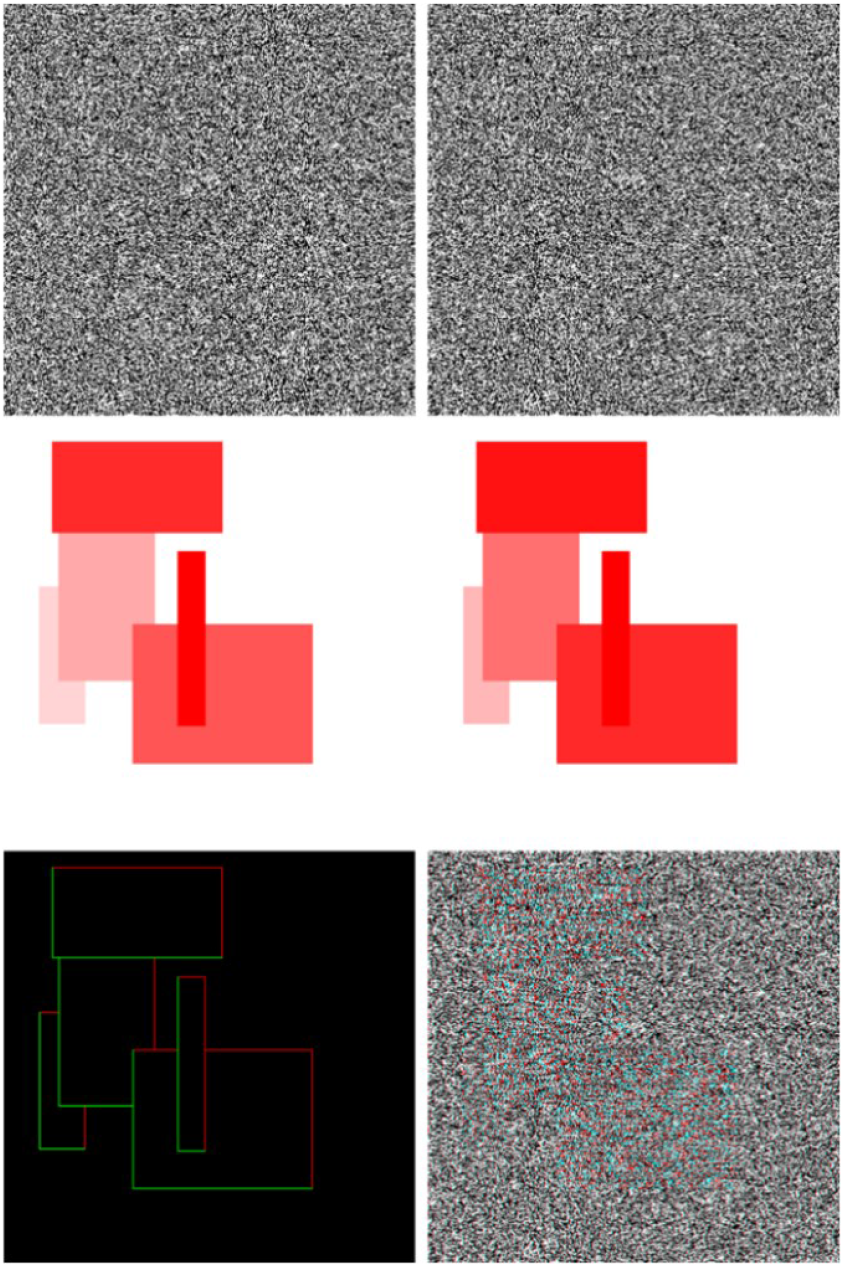
Random Dot Stereograms dataset (adapted from Fig. 8 in Chen et al., 2025; dataset from Chen, 2025a). **Top**: Left and right RDS images depict disparity-defined objects created by random dot shifts. **Middle**: Near and Far disparity maps (red intensity inversely proportional to depth). **Bottom**: 2-channel OC border-ownership maps for disparity-defined objects and the corresponding stereo anaglyph.

**Figure 6.**
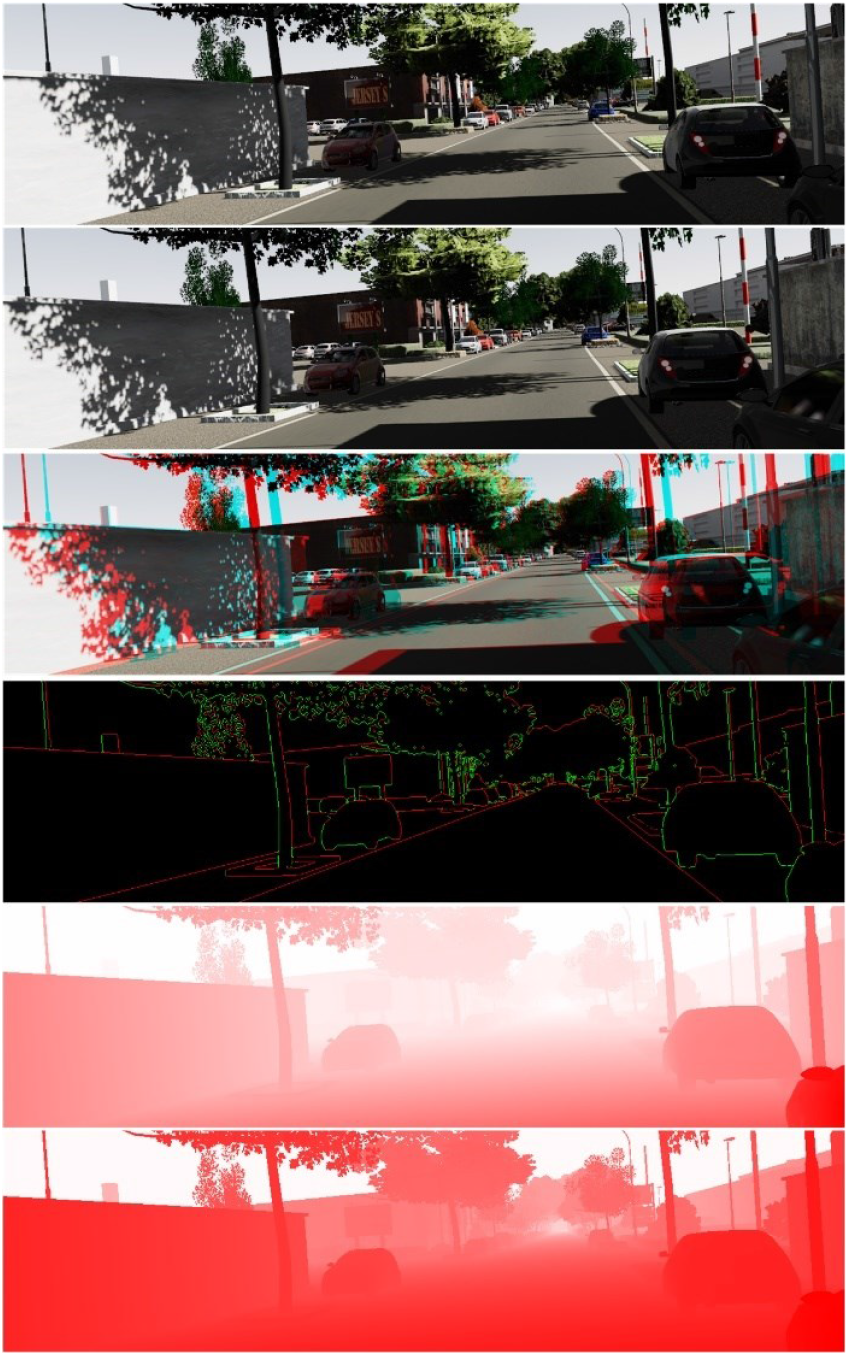
Modified Virtual Kitti 2 Dataset (adapted from Fig. 5 in Chen et al., 2025; dataset from Chen, 2025b; original VKitti from Cabon et al., 2020). **From top to bottom**: stereo left and right images, stereo anaglyph, 2-channel border-ownership OC coding map (left-view), Near (Epipolar) disparity map (darker red corresponds to closer depths), and Far (Logarithmic) disparity map capturing depth order across large distances.

**Figure 7.**
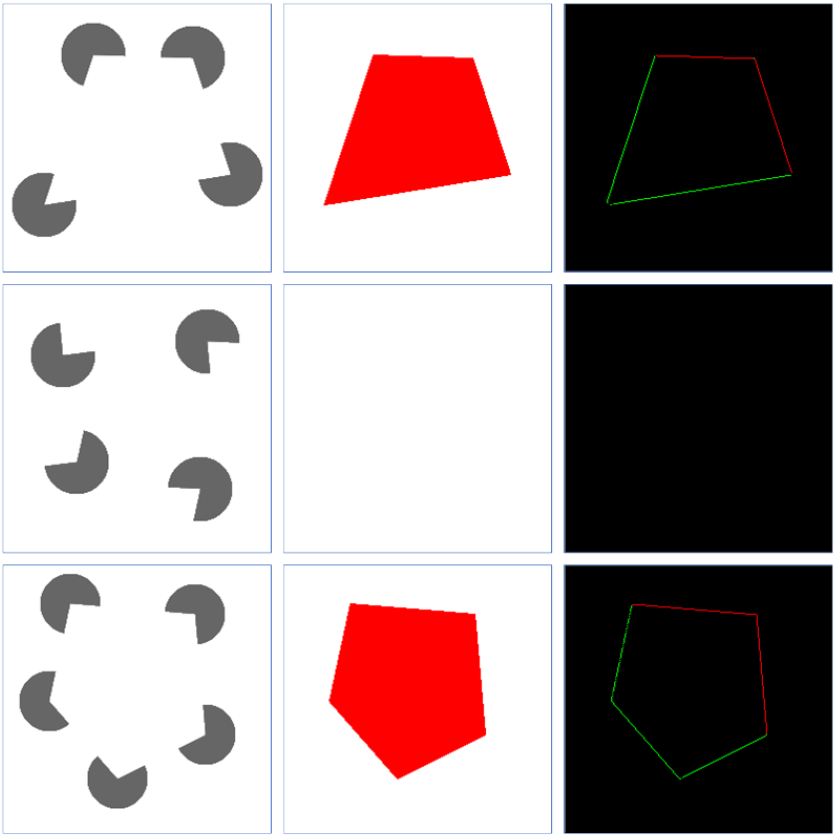
Kanizsa figures with synthetic Near disparity. (adapted from Fig. 9 in Chen et al., 2025; dataset from Chen 2026). **From left to right**: Kanizsa figures, synthetic Near disparity maps (disparity=2 pixels), and 2-channel OC border-ownership maps. The **middle row** shows “false” Kanizsa figures created by rotating endpoints by 90° to disrupt illusory closure.

Together, these datasets span the three major domains of figure–ground organization: disparity-defined (RDS), contrast-defined (VKitti), and illusory-defined (Kanizsa). This combination provides a comprehensive testbed for evaluating whether RD coding serves as a generalizable depth-driven substrate that achieves performance comparable to OC coding across physical, photorealistic, and illusory contexts.

### 5.3. Experimental Setup

#### 2-Channel OC vs RD Coding

The experiments were designed to directly compare opposite-channel-rule (OC) coding and relative-disparity-based (RD) coding under controlled conditions. Unlike our previous work (Chen et al., 2025), the present experiments used absolute disparity maps directly— either ground truth (perfect) or intermediate disparity representations—without stacked RGB image inputs.

This isolates the role of disparity itself, enabling a clean evaluation of the extent to which disparity-driven mechanisms alone can account for figure–ground organization, and whether thresholded relative-disparity signals can replicate or approximate the border-ownership patterns produced by OC-trained TcNet.

A *training-free* TcRd model was implemented, taking absolute disparity as input and applying relative-disparity (RD) sign coding (Figure 4(b)). Relative disparity was computed following (Eq. 4.7), and the resulting thresholded RD maps were compared directly with corresponding TcNet outputs generated using OC coding. For completeness, the ‘all-contour’ channel (also referred to as the ‘enforcement’ channel in Chen et al., 2022) was also generated by summing the two border ownership channels.

This setup enables side-by-side evaluation of rule-based (OC) versus disparity-based (RD) coding under equivalent conditions, as illustrated in Figure 8.

**Figure 8.**
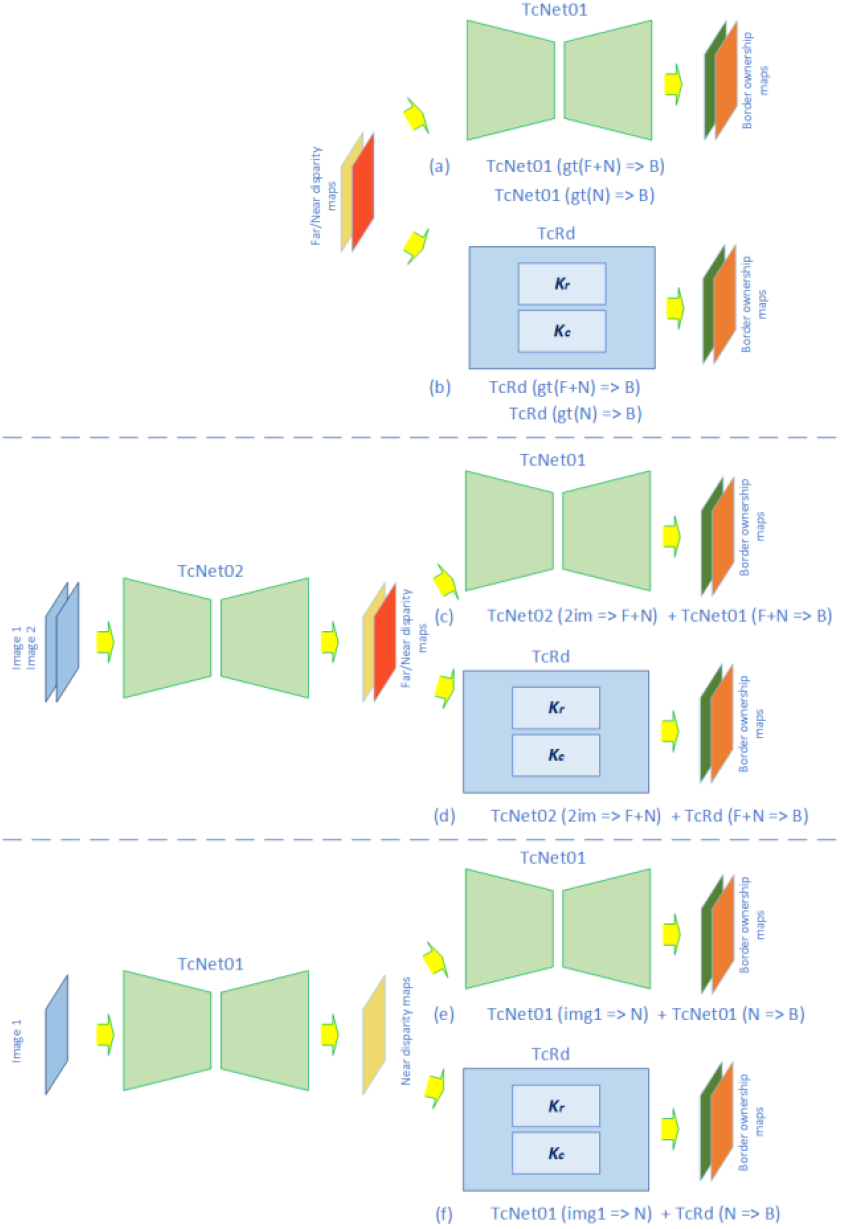
Three TcNet vs TcRd Comparison Setups. **(a, b)** Ground truth Far/Near disparities used directly as input to generate border ownership maps (perfect-disparity condition). **(c, d)** Far/Near disparities first computed as intermediate representations from stereo image pairs, then used to generate border ownership maps. **(e, f)** Near disparity computed as intermediate representation from a single image, then used for border-ownership generation.

#### 2-Channel vs 1-Channel Coding

To test whether the *sign of relative-disparity alone* can serve as a viable basis for opposite owner-side coding, the second channel of a 2-channel OC coding was negated and added to the first channel to form a 1-channel representation.

This 1-channel configuration was then compared with its 2-channel counterpart to assess whether sign-based opposite coding is sufficient to capture the correct owner-side selectivity.

#### Methodology Consistency

To ensure methodological consistency, all network configurations followed the **same notation conventions** of Chen et al. (2025). The following additional notations were introduced for clarity:

- A numerical suffix was added to indicate the number of border-ownership channels: for instance, **B2** and **B1** denote 2-channel and 1-channel border-ownership coding, respectively. **B2(RD)** indicates RD based sign coding, otherwise B2 denotes OC based coding.
- **+C** denotes inclusion of category-selective channels, so **B+C** denotes the joint configuration of border-ownership and category-selectivity outputs.
- **gt(F+N)** and **gt(N)** indicate ground-truth (perfect) *Far + Near* and *Near* disparity inputs, respectively, rather than intermediate disparities derived from earlier processing stages (Figure 8(a-b)).
- **TcRd(k1.5)** and **TcRd(k3.4)** specify that the corresponding TcRd implementations employed **k1.5** and **k3.4** kernel sets, respectively.

These notations are applied consistently throughout the paper to allow unambiguous comparison across datasets and model configurations, ensuring clarity when interpreting the results of OC- and RD-based coding schemes.

#### Red-Green Display Coding Scheme

Border-ownership visualizations follow the same red-green scheme used in our prior work (Chen et al., 2022; Chen et al., 2025) and as illustrated in Figure 5 (Bottom left).

In this scheme, red segments indicate borders whose owner lies on the ‘left’ or ‘bottom’ side, while green segments indicate borders whose owner lies on the ‘right’ or ‘above’ side.

### 5.4. Statistical Evaluation Criteria

The statistical evaluation criteria used in this study follow exactly the procedures described in Chen et al., (2025), ensuring consistent and fair comparison across all configurations. As in that work, only the final outputs— specifically, the contour maps and border-ownership maps—were statistically evaluated, while intermediate results (including disparity maps) were excluded from evaluation.

#### (a) Mean Average Precision (mAP)

Mean average precision (*mAP*) was calculated by averaging the ratio of correctly predicted contour points over all predicted contour points across all test cases, specifically,

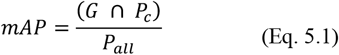

where *G* denotes ground truth, *P*_*c*_ and *P*_*all*_ represent, respectively, correct prediction and all prediction (both correct and incorrect).

#### (b) Thresholding and Contour Thinning

A threshold was applied to the results, and all contours were thinned to one-pixel width before statistical calculations. However, the threshold values differ between coding schemes due to their distinct computational structures:

- **OC coding (TcNet):** threshold > 0.1
- **RD coding (TcRd):** threshold > 0.001

*The threshold values were empirically selected to be balanced*—sufficiently high to suppress noise yet low enough to retain valid contour detections.

#### (c) 1-Neighbor Ground-Truth Criterion

A 1-neighbor criterion was used when comparing predicted contours with ground truth: a predicted contour point was counted as correct only if it fell within one pixel of a ground-truth contour point.

#### (d) ‘False’ (Empty) Case Criterion

For ‘false’ (empty) cases, evaluation was based on the accuracy of predicting empty maps. A predicted map was counted as *empty* only if it contained no more than 36 active pixels, corresponding to at most four false endpoints in an empty Kanizsa figure (see Figure 7; 4 end-points × 9 pixels each = 36).

Accuracy was computed as:

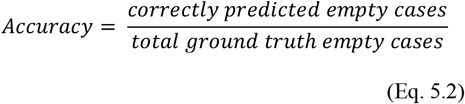

### 5.5. Results

#### 5.5.1. Quantitative Evaluation

To ensure consistency and enable direct comparison across datasets, all results were evaluated using the same statistical criteria described in Section 5.4.

For reference, the first several rows in each results table (Table 1 for RDS, Table 2 for VKitti, and Table 3-Table 4 for Kanizsa) are taken directly from Chen et al. (2025) to serve as benchmarks.

**Table 1.**
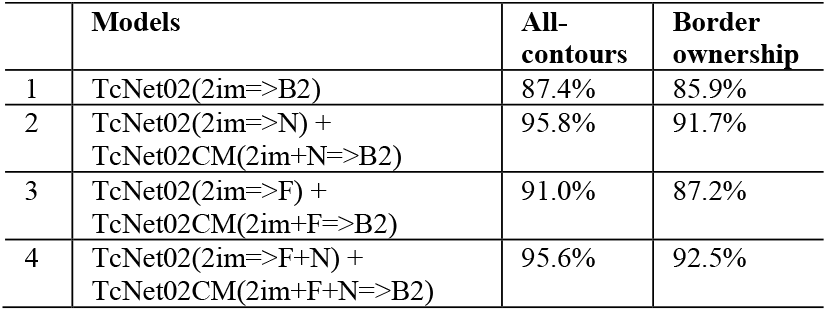
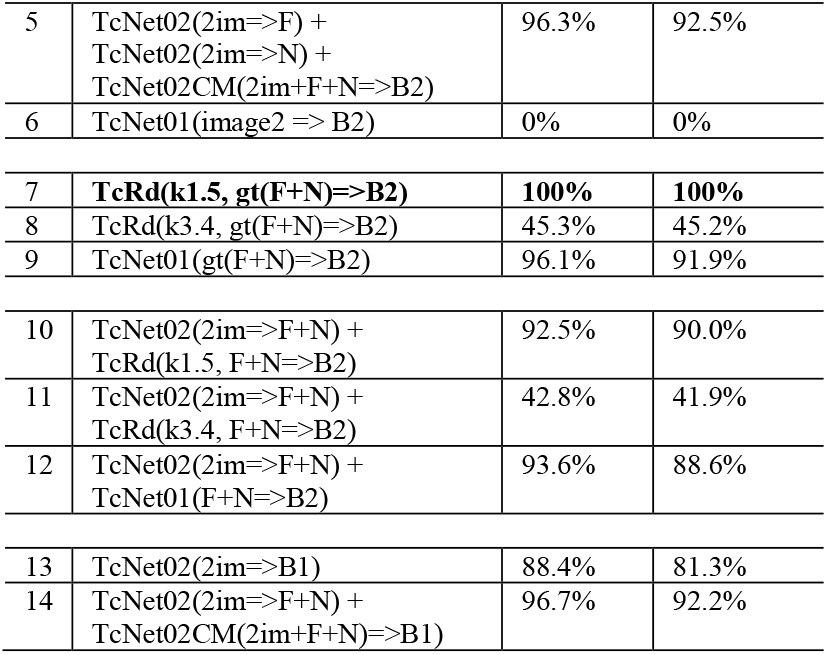
Evaluation results for the Random Dot Stereograms (RDS) dataset. Rows 1–6 are taken directly from Chen et al. (2025, Table 1). Subsequent rows report the new evaluation results for the OC and RD coding schemes on the same dataset.

**Table 2.**
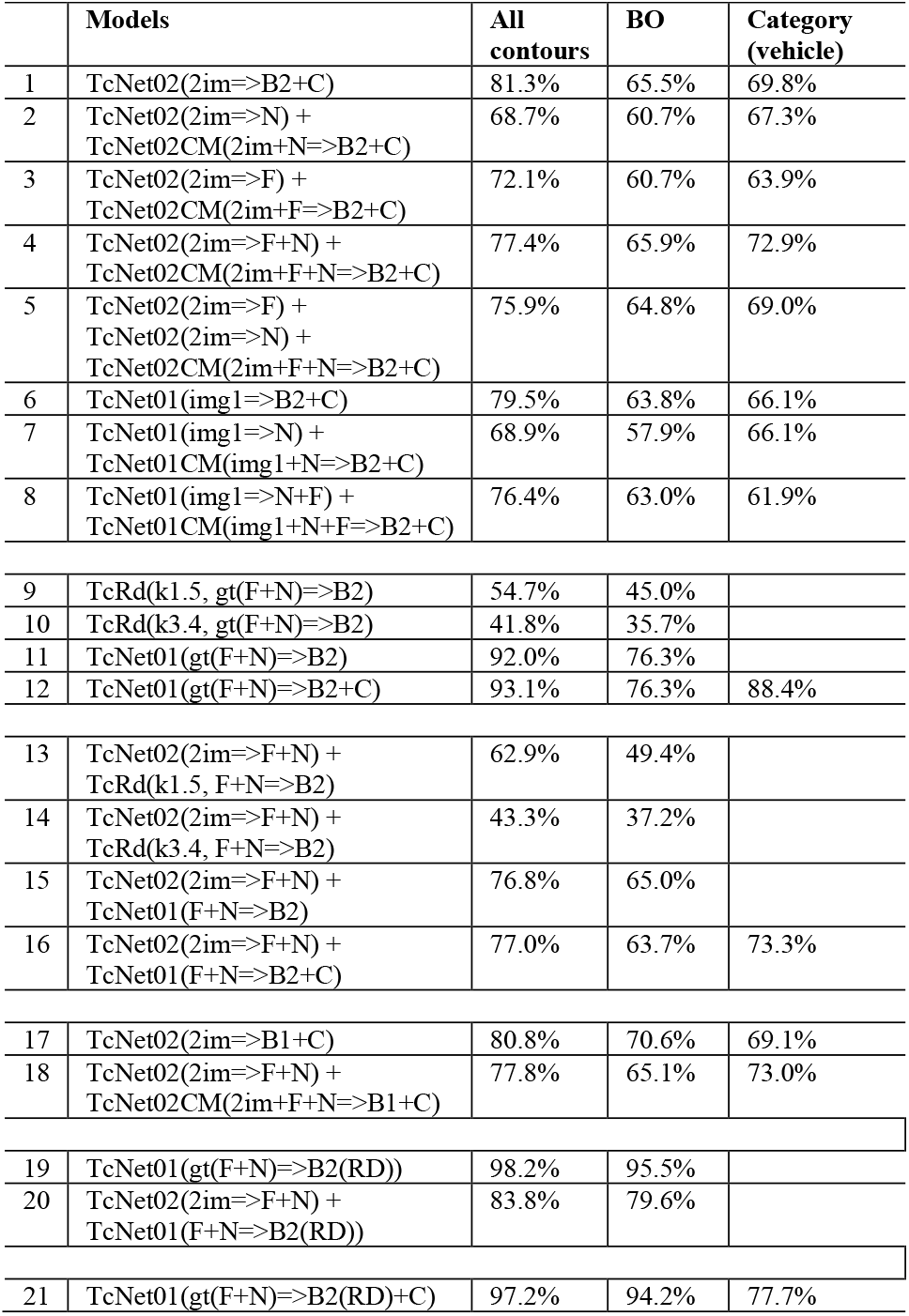
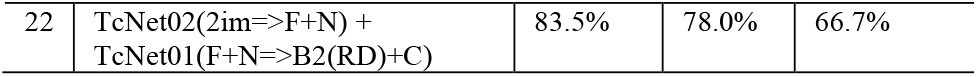
Evaluation results for the modified Virtual Kitti 2 (VKitti) dataset. Rows 1–8 are taken directly from Chen et al. (2025, Table 2). Subsequent rows report the new evaluation results for the OC and RD coding schemes on the VKitti dataset.

**Table 3.** Monocular evaluation results for Kanizsa Figures with synthetic Near disparity (disparity=2). Groups (1) and (2) are taken directly from Chen et al. (2025, Tables 3 and 6, respectively). Group (1) corresponds to Kanizsa figures without synthetic Near disparity. Subsequent rows report the new evaluation results for the OC and RD coding schemes under monocular conditions.

|  | All-contours | Border ownership |
| --- | --- | --- |
| (1) TcNet01(img1=>B2) |  |  |
| Triangle/quadrilateral | 97.7% | 91.6% |
| 'false' (empty) | 96.4% |  |
| Pentagon | 65.5% | 60.3% |
| Hexagon | 75.9% | 72.8% |
| (2) TcNet01(img1=>N) + TcNet01CM(im1+N=>B2) |  |  |
| Triangle/quadrilateral | 73.7% | 50.0% |
| 'false' (empty) | 0% |  |
| Pentagon | 37.9% | 22.7% |
| Hexagon | 43.0% | 26.5% |
| (3) TcNet01(img1=>N) + TcNet01(N=>B2) |  |  |
| Triangle/quadrilateral | 67.8% | 33.7% |
| 'false' (empty) | 0% |  |
| Pentagon | 38.4% | 17.7% |
| Hexagon | 33.0% | 14.9% |
| (4) TcRd(k1.5, gt(N)=>B2) |  |  |
| Triangle/quadrilateral | 97.7% | 84.8% |
| 'false' (empty) | 100% |  |
| Pentagon | 95.9% | 80.9% |
| Hexagon | 93.3% | 85.9% |
| (5) TcNet01(img1=>N) + TcRd(k1.5, N=>B2) |  |  |
| Triangle/quadrilateral | 76.9% | 69.9% |
| 'false' (empty) | 0% |  |
| Pentagon | 38.3% | 31.4% |
| Hexagon | 41.9% | 36.8% |
| (6) TcNet01(gt(N)=>B2) |  |  |
| Triangle/quadrilateral | 93.2% | 84.9% |
| 'false' (empty) | 100% |  |
| Pentagon | 83.7% | 72.2% |
| Hexagon | 56.2% | 48.9% |

**Table 4.**
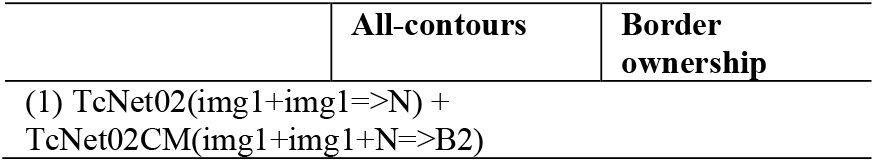
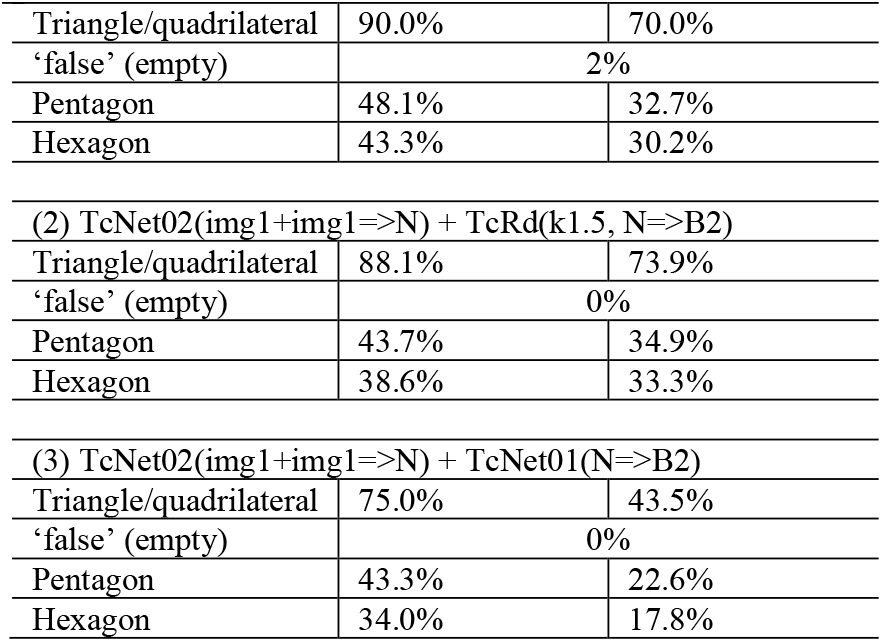
Binocular evaluation results for Kanizsa Figures with synthetic Near disparity (disparity=2). The (1) group is taken directly from Chen et al. (2025, Table 7). Subsequent rows report the new evaluation results for the OC and RD coding schemes under binocular conditions.

The remaining rows report new evaluations using the OC and RD coding schemes applied under their corresponding test configurations, including both 2-channel and 1-channel variants.

Key findings from these quantitative results are summarized in Section 5.5.2 (Key Observations).

#### 5.5.2. Key Observations

The following observations synthesize the main empirical findings drawn from the quantitative results in Tables 1–4, highlighting patterns that clarify the relationship between relative-disparity-based (RD) and opposite-channel-rule (OC) coding schemes.

##### (a) Kernel and Computational Equivalence

The ***most striking***—and *theoretically anticipated*— result appears in Row 7 of Table 1 (**RDS**), where the *TcRd(k1*.*5)* model with *RD sign coding* produces outputs ***identical (100%)*** to those generated by the OC-based configuration on the RDS dataset when supplied with perfect absolute (Far/Near) disparity input (Figure 8 (a-b)).

This provides **direct empirical confirmation** that, under an appropriate kernel structure, ***thresholded relative disparity differencing can reproduce the functional mapping of border ownership*** generated by rule-based OC coding.

In contrast, *TcRd(k3*.*4)* performs substantially worse than *TcRd(k1*.*5)* across both RDS and VKitti (Rows 8 **≪** 7 and 11 **≪** 10 in Table 1; Rows 10 ≪ 9 and 14 ≪ 13 in Table 2), establishing that:

- *kernel structure is critical*;
- ***k1*.*5 closely matches the mathematical definition of relative disparity***; and
- k3.4 integrates disparity differences in a manner misaligned with OC-style owner-side assignment.

The performance gap between k1.5 and k3.4 aligns with the broader distinction discussed in Section 6.3.2, where these operators exhibit different behaviors in long-range relative disparity.

##### (b) Disparity Quality (Perfect vs Intermediate)

The equivalence between RD- and OC-based representations persists even when disparity inputs are not perfect.

For the **RDS** dataset using intermediate (Far/Near) disparity, TcRd(k1.5) achieves performance comparable to OC-based TcNet (Row 10 ≈ Row 12 in Table 1; Figure 8(c-d)), indicating that local disparity differencing remains effective with non-ideal disparity.

In contrast, for the photorealistic **VKitti** dataset with perfect disparity input (Row 9 in Table 2), *TcRd(k1*.*5)* underperforms relative to OC-trained TcNet01 (Rows 9 **≪** 11 or 12).

Visual inspection—comparing the ground-truth border-ownership map (Figure 6 Row 4) and disparity maps (Figure 6 Rows 5–6) with the TcRd(k1.5) output (Figure 9 Row 3)—reveals three contributing factors:

**Figure 9.**
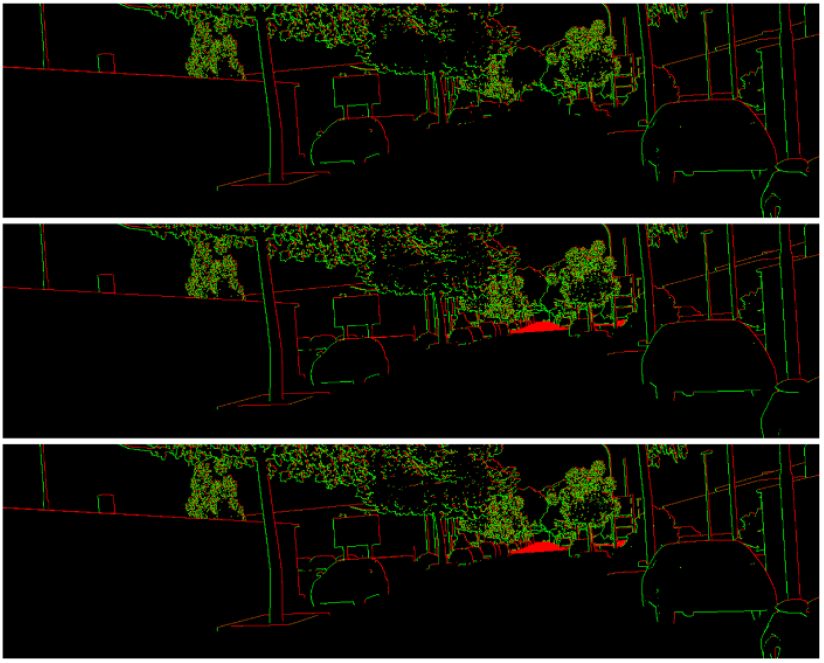
TcRd(k1.5) performance on a sample from the modified Virtual Kitti 2 (VKitti) dataset (sample shown in Figure 6; dataset from Chen, 2025b; original VKitti from Cabon et al., 2020). **From top to bottom**: RD based border ownership maps from Near, Far and Sum (= Near + Far) disparities. The summed map visualizes the integrated ownership signals from both disparity layers, providing a *unified representation* across depth channels within the *layered disparity framework*.

- several annotated contours in VKitti correspond to subjective or perceptual edges (e.g., curb lines) that lack disparity support;
- some genuine disparity-defined transitions (e.g., road-wall boundaries) are missing from the ground-truth BO labels; and
- RD-based maps allow row/column channel overlap, which is penalized during OC-based evaluation because OC coding assumes exclusive, non-overlapping owner-side channels.

Thus, the discrepancy reflects a *mismatch* between disparity structure and OC-based ground truth— combined with *overlap penalization*—rather than any inherent limitation of the RD differencing operation.

To validate this interpretation, we performed additional experiments in which:

1. TcRd(k1.5) was used to generate ground truth RD-based border ownership (with the category-selective channel was taken directly from OC-based ground truth to form a **B2+C** reference), and
2. TcNet01 was then trained to predict BO from either perfect (ground-truth) or intermediate disparity inputs (Rows 19-22, Table 2).

The results confirmed the expected pattern:

- Perfect disparity → performance approached 100% (Row 19 ≈ Row 21 ≈ 95% ≫ Row 11 for BO);
- Intermediate disparity → performance remained significantly higher than the corresponding OC-based baseline (Row 20 ≈ Row 22 ≫ Row 15).

These findings verify that *RD-based ownership becomes nearly indistinguishable from OC-based ownership when the disparity–ownership structure is internally consistent*.

They further demonstrate that *TcNet can be trained to emulate TcRd(k1*.*5)*, supporting the view that *disparity differencing alone is a sufficient and biologically interpretable basis* for border-ownership generation.

##### (c) Sign-Based Opposite Coding

Across both databases—**RDS** (Row 13 ≈ Row 1 and Row 14 ≈ Row 4 in Table 1), and **VKitti** (Row 17 ≈ Row 1 and Row 18 ≈ Row 4 in Table 2)—the 1-channel coding configurations exhibit performance comparable to their 2-channel counterparts.

These results indicate that:

- *the sign of relative disparity alone is sufficient for owner-side selectivity*.
- ***sign-based opposite coding*** *appears to function as* ***a general computational mechanism*** that *plausibly underlies* both relative disparity and border ownership representations.

##### (d) Dataset-specific Geometry and Directional Simplicity

The **RDS** dataset is unique among the tested datasets in that, although its channel structure *theoretically* allows horizontal–vertical overlap, the stimulus geometry ensures that the horizontal and vertical components remain *strictly non-overlapping* in practice (Figure 3 and Figure 5).

This geometric simplicity makes RDS an ideal testbed for assessing the separability of directional components in border-ownership computation. Under these conditions, both OC- and RD-based coding schemes—across 1-, 2-, and 4-channel configurations—produce border-ownership maps that are *identical* (100% correspondence).

This outcome reflects the *inherent alignment* between disparity structure and owner-side assignment in RDS.

##### (e) Disparity with and without Stacked RGB

In our prior work (Chen et al., 2025), stacked RGB channels were included as part of the intermediate representation to provide additional contextual information—implicitly capturing cues beyond pure disparity—during border-ownership generation.

For the **RDS** dataset, the TcNet configuration using intermediate disparity input stacked with RGB outperforms both the intermediate disparity input without RGB and the direct RGB input alone (Row 4 > Row 12 > Row 1 in Table 1).

This outcome aligns with the fact that *RDS represents a depth-only, weak-context case*, and the inclusion of additional RGB provide supplementary contextual cues that are absent from pure disparity signals.

In contrast, for the **VKitti** dataset, the TcNet configuration using intermediate disparity stacked with RGB yields comparable performance to both disparity-only and RGB-only configurations (Row 4 ≈ Row 16 ≈ Row 1 in Table 2).

This suggests that, in photorealistic scenes, the *intermediate disparity representation already encodes rich contextual structure*—implicitly learned from the underlying scene geometry—rendering the addition of RGB largely redundant. In such cases, *either* stacked RGB or layered disparity alone is sufficient.

(see further discussion in Section 6.3.8)

##### (f) Illusory Disparity

We revisit the **illusory Kanizsa figures with synthetic Near disparity** (Chen et al., 2025). Across both monocular configurations (Groups (2), (3), and (5) in Table 3) and binocular settings (Table 4), whether using TcNet01 or TcRd(k1.5) to process the intermediate disparity, performance is markedly degraded relative to the direct generation of border-ownership maps from the Kanizsa figures themselves (Group (1) in Table 3).

Both the benchmark statistics and qualitative inspection indicate that this degradation is driven primarily by the quality of the intermediate disparity extracted from the illusory stimuli. The generated disparity maps contain distortions or inconsistencies that do not align with the true illusory structure, thereby limiting subsequent computation of border ownership.

This interpretation is strongly supported by two additional experiments in which TcRd(k1.5) (Group (4) in Table 3) or TcNet01 (Group (6) in Table 3) were supplied with perfect ground-truth disparity to generate border-ownership maps. In both cases, performance became comparable—judged by the first row in each group—to the direct border-ownership generation obtained by TcNet01 (Group (1) in Table 3).

These findings indicate that the performance degradation stems from limitations in disparity extraction rather than any weakness of the RD/BO mechanism itself. The implications for the representation of illusory depth representation are examined further in Section 6.3.1.

## 6. Integrative Discussion

### 6.1. Re-interpretation of Border Ownership as Thresholded Relative Disparity

The results presented in this study support a major reinterpretation of ***border ownership (BO)***: what has been regarded as a distinct neural mechanism for assigning figure–ground directionality can instead be understood as an ***emergent consequence of thresholded relative disparity***. In this view, border-ownership–selective responses arise when local disparity differences exceed a threshold, and the ***sign of the directed disparity difference determines the owner-side direction***.

In other words, the ***biological basis of border-owner selectivity may derive directly from the magnitude and sign of the local directed disparity difference***, rather than from an independently specialized BO computation.

### 6.2. Functional Implications of This Re-interpretation

#### 6.2.1 Unification of BO and RD

If border ownership (BO) arises from thresholded relative disparity (RD), then BO and RD are not separate visual processes but two expressions of the same underlying computation—a local ***directed-disparity-differencing operation*** *followed by* ***thresholding***.

Under this interpretation, ***depth ordering and figure– ground segmentation become intrinsically linked***: *both emerge from a single, unified, and biologically grounded mechanism for early visual organization*.

#### 6.2.2 A Mechanistically Transparent Explanation of Classical BO Findings

A thresholded relative-disparity (RD) model—where border ownership (BO) arises from local directed disparity differences followed by thresholding—naturally clarifies several long-standing findings in visual neuroscience.

Under this interpretation, many classical neurophysiological findings become immediate and mechanistically self-explanatory:

##### (a) Why border-ownership neurons are sensitive to depth edges and figure-ground relation

V2 neurons exhibit owner-side selectivity that aligns with depth order: “*the ‘near’ side of the preferred 3D edge generally coincides with the preferred side-of-figure in 2D displays*” (Zhou et al., 2000; Qiu & von der Heydt, 2005)

Under the RD interpretation, this correspondence emerges automatically because *owner-side direction is determined by the sign of the directed disparity difference*.

##### (b) Why relative disparity yields more stable perceptual judgements than absolute disparity

Relative disparity is fixation-invariant because it depends only on the *difference* between disparities, not absolute values.

If BO is a directed form of RD, then BO inherits this invariance: whenever local depth relations are preserved, the owner-side direction remains stable across fixation changes and global positional shifts.

This provides a simple mechanistic account for the well-known stability of both RD and BO across eye movements and viewing conditions.

##### (c) Why both border-ownership and relative-disparity signals first appear in V2

Both computations require *pooling* from multiple V1 disparity-tuned or orientation-tuned units:

- relative disparity requires integrating across spatially separated V1 inputs (Tanabe et al., 2008; Thomas et al., 2002);
- border ownership requires integration across oriented inputs spanning a contour (Zhou et al., 2000).

Directed disparity differencing requires exactly this V2-scale pooling, explaining why both RD and BO first arise in V2 with similar latencies and spatial integration characteristics.

##### (d) Why border ownership arises with short latency

Border-ownership signals arise at short latency (∼25 ms) in V2 (Zhou et al., 2000) reflecting the minimal feedforward nature of the computation: a simple local differencing followed by thresholding.

##### (e) Why border ownership is invariant to cue, size and position

Because relative disparity–based border ownership depends on local depth relations rather than on the specific visual cues that give rise to a border, BO selectivity is invariant across different inducing cues, spatial scales, and positions.

Consistent with this, neurophysiological studies from the same research program report that “*many cells showed invariance to the cues that produce perception of border ownership*” and exhibited *size- and position-invariant* responses (Zhou et al., 2000; Craft et al., 2007).

##### (f) Why owner-side direction is orthogonal to the border

Because the directed disparity difference is computed *across* the border (Figure 3), the resulting owner-side direction naturally lies orthogonal to it. This is consistent with neurophysiological observations that the border-ownership signal is “position invariant in the direction orthogonal to the border” (Williford et al., 2013). This provides a *direct mechanistic explanation of owner-side direction* (see further discussion in Section 6.3.3).

##### (g) Why border-ownership signals remain stable across saccades, motion, and brief occlusion/disocclusion events

Border-ownership tuning remains stable even when a contour temporarily disappears—during a saccade, object motion, or brief occlusion—and later reappears. Newly activated neurons inherit the previous owner-side assignment (O’Herron et al., 2013; see also Zhou et al., 2000).

Under the thresholded-RD model, this stability follows naturally: the computation depends on local directed disparity differences, not on absolute retinal position or momentary visibility. Once the same local disparity relations are restored after the shift or occlusion, the same owner-side direction re-emerges automatically.

Thus, persistence across saccades, motion, and brief occlusion is an inherent consequence of the *relational (difference-based)* nature of RD-based border ownership.

##### (h) Why border ownership reverses under depth reversal

Border-ownership responses reverse polarity when the perceived depth order of two regions reverses (Zhou et al., 2000; Qiu & von der Heydt, 2005).

In the thresholded-RD framework, this is immediate: reversing depth order simply reverses the sign of the directed disparity difference. The owner-side direction therefore flips automatically, with no need for an additional mechanism or reinterpretation.

Thus, depth reversal entails a straightforward algebraic sign change in the directed disparity signal, yielding a direct and mechanistically transparent account of polarity reversal.

Together, these observations converge on **a single principle**: *border ownership and relative disparity reflect a unified, feedforward computation based on thresholded disparity differencing*.

This shared mechanism provides a simple and mechanistically transparent explanation for the common latency, cortical locus in V2, perceptual stability across saccades and motion, and automatic polarity reversal under depth-order changes observed in both BO and RD.

Moreover, numerous additional properties of BO— such as its consistency across diverse stimulus classes and robustness under contextual manipulation—arise naturally from the relational structure of the RD computation, reinforcing the interpretation that BO is not an independent process but a directed expression of relative disparity.

#### 6.2.3 Confirmation and Strengthening of Our Prior Assertion

The thresholded-RD interpretation provides a coherent foundation for several key assertions made in our earlier work (Chen et al., 2025), which now emerge as natural consequences of the intrinsic disparity-differencing computation:

##### (a) BO generation is intrinsic and concurrent with border formation

Because the ***border itself*** arises from thresholded directed disparity differencing, the ***owner-side direction*** is determined at the same moment that the border becomes encoded.

Thus, BO emerges concurrently with border formation—across contrast-defined, disparity-defined, illusory, and contour-defined boundaries—supporting our earlier definition of an *object border* as any contour, visible or illusory, that carries an ownership state encoding relational structure.

##### (b) T-junctions are not causal cues

Under RD model, BO is not “assigned” post-hoc based on symbolic cues but emerges directly from local disparity differences.

This clarifies why T-junctions correlate with BO but do not generate it causally.

##### (c) Binocular BO is not derived from fusing left/right monocular borders

In the RD framework, border ownership is generated directly from the signed binocular disparity, not from combining or matching monocular borders. Thus, BO arises intrinsically from disparity differencing rather than through any contour-fusion process.

##### (d) Consistency with feedforward surface filling-in along owner-side direction

The orthogonal encoding of the derived owner-side direction is consistent with our previously proposed feedforward surface-filling-in model, in which ownership signals propagate preferentially along the owner-side direction to support *rapid surface completion and region formation*.

Once the owner-side direction is encoded at the border, this direction—corresponding to the direction of increasing disparity—is *naturally utilized* as the propagation direction, allowing surface filling-in to proceed without additional contour-based routing. The present RD-based formulation therefore provides a mechanistic origin for the owner-side propagation assumed in the earlier model.

##### (e) The k1.5 operator aligns with “active neuron” locality

The k1.5 kernel implements a strictly *local* differencing operation, consistent with our earlier proposal that BO-selective “active neurons” act as *local*, semi-independent units that coordinate to achieve global figure–ground organization.

##### (f) BO and disparity are tightly coupled within a unified computation

The RD framework formalizes the tight coupling between depth ordering and figure–ground assignment, reinforcing the integrative basis of the active-neuron framework (Chen et al., 2025), in which local units are dynamically linked along ownership contours to maintain object coherence.

Taken together, these points show that the thresholded– RD interpretation not only supports but also strengthens the ***core principles*** underlying our earlier framework: border ownership arises intrinsically, operates through strictly local computations, and is fundamentally tied to disparity structure rather than symbolic contour cues.

This convergence provides a *coherent conceptual foundation* for the active-neuron model and sets the stage for the deeper mechanistic analyses developed in Section 6.3 and the V4-mediated feedback mechanism examined in Section 6.4.

### 6.3. Nature and Structure of Disparity

Having established the intrinsic computational equivalence between border ownership and thresholded relative disparity, we now examine the structure and functional organization of disparity and of the derived representational signals, including border ownership and category-selective channels.

#### 6.3.1 Epipolar and Non-Epipolar Disparity

The empirical results summarized in Section 5.5.2(f) show a clear and highly diagnostic pattern: border-ownership (BO) performance degrades *only* when disparity must be *estimated* from Kanizsa inputs, but is fully restored when either perfect ground-truth disparity is supplied (Groups (4) and (6) in Table 3), or BO is generated directly from the Kanizsa figure itself (Group (1)).

This pattern indicates that the performance collapse arises *not* from any limitation of the BO mechanism, *but rather* from failures in the *disparity-estimation* stage under illusory conditions.

Two explanations were possible when we revisited this phenomenon (see also Section 3.4):

##### (1) Attentional-highlighting hypothesis (Chen et al., 2025)

The perceived depth in Kanizsa figures might be generated by a process entirely separate from BO—for example, an attentional-highlighting enhancement— implying that illusory depth and BO are produced independently.

##### (2) Different-disparity-type hypothesis

The depth perceived in Kanizsa configurations might arise from a *qualitatively different form of disparity* that does not obey epipolar geometry and is therefore poorly captured by TcNet’s disparity estimators, which were designed for physically grounded (epipolar) disparities in RDS and VKitti scenes.

If the hypothesis that thresholded relative disparity represents the neural basis of border ownership is correct, then only the second explanation remains viable—namely, that ***relative disparity (and consequently border ownership) must always originate from some form of disparity representation, whether physically or perceptually constructed***.

This motivates a principled distinction between *epipolar* and *non-epipolar* (or *illusory*) disparity processes.

#### Epipolar Disparity

The absolute disparities represented in the RDS and VKitti datasets linearly or non-linearly follow epipolar geometry (Chen et al., 2025) and are therefore referred to here as *spatial epipolar disparities*.

In contrast, motion parallax arises when relative motion occurs between the observer and the surrounding scene— either through observer movement or object motion— causing the same spatial point to be sampled from slightly different viewpoints over time. This motion creates an *effective temporal baseline*—the path traced by the observer’s translation relative to the scene—such that the geometry of motion parallax becomes analogous to the binocular baseline. Accordingly, the depth cue derived from optic flow can be regarded as a form of *temporal epipolar disparity*.

Spatial and temporal epipolar disparities both originate from physical 3D scenes and may be collectively described as *physical epipolar disparities*.

In addition, when an observer “looks into” a 2D image, a virtual or perceptual 3D scene can be experienced that implicitly follows epipolar geometry; the perceived disparity in such cases is thus referred to as *perceptual epipolar disparity*.

#### Illusory (Non-Epipolar) Disparity

By contrast, the disparity perceived in illusory configurations such as Kanizsa figures appears not to follow epipolar geometry and is therefore referred to here as *non-epipolar disparity*, or simply *illusory disparity*.

A hypothetical subtype, referred to as *Category Illusory Disparity* (see Section 6.3.5), may arise when the inferred “disparity” is guided primarily by categorical or semantic context rather than being strictly constrained by local geometric cues.

This hypothesis suggests that category-consistent or other high-level priors could give rise to illusory disparity components that bias figure-ground interpretation, thereby linking categorical or contextual information with *disparity-based perceptual organization*.

Accordingly, illusory disparity may be viewed more broadly as a form of *contextual disparity* that draws on geometric cues but is primarily shaped by *higher-level contextual priors* acting together with non-epipolar geometric regularities.

#### Spectrum of Disparity Forms

The distinctions introduced above—between epipolar, perceptual, and illusory (non-epipolar) disparities— provide a conceptual framework for describing multiple depth signals. Empirical and perceptual evidence further shows that these different forms of disparity can coexist and interact within the visual system.

Figure 10 illustrates this variety: the same 2-D image can yield different perceptual outcomes depending on contextual interpretation and perceptual grouping.

**Figure 10.**
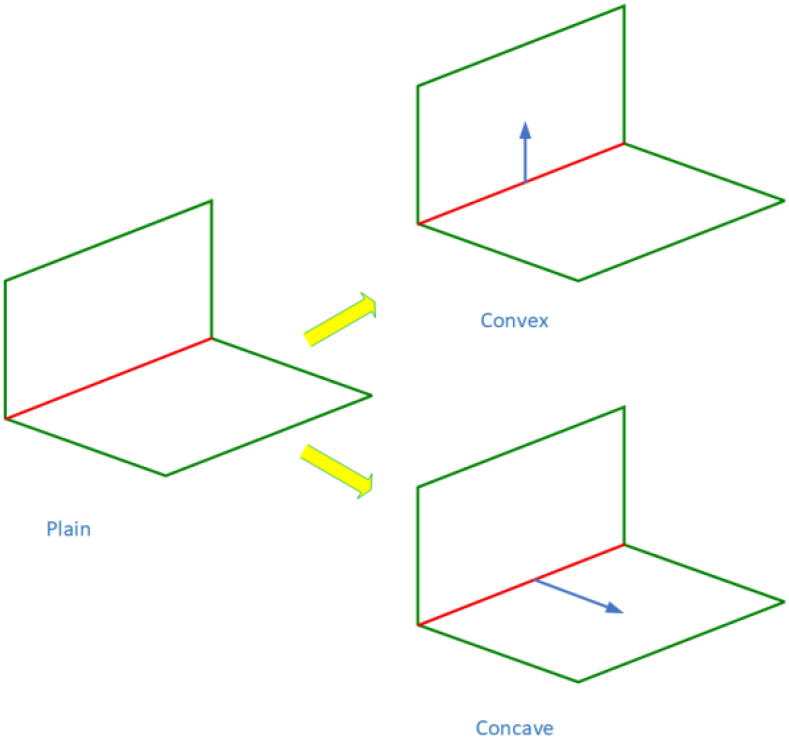
Examples of Illusory Disparity and Perceptual Epipolar Disparity. The same 2D image can be perceived in multiple ways. **Left**: An illusory disparity perceived from a plain 2D figure, where depth is inferred without any epipolar constraint and the red segment does not correspond to an actual object border; alternatively, no disparity is perceived when no distinct object is recognized. **Right (top)**: A perceptual epipolar disparity perceived when the vertical rectangle appears in front of a horizontal, lay-down rectangle—the red border belongs to the vertical object. **Right (bottom):** A perceptual epipolar disparity perceived when the horizontal, lay-down rectangle appears in front of a vertical rectangle—the red border belongs to the horizontal object.

- When no distinct object is recognized, no disparity is experienced.
- When a surface or figure emerges without geometric constraint, *illusory disparity* arises, reflecting depth inferred primarily from contextual and semantic priors.
- When the same image is interpreted as a coherent 3-D configuration consistent with binocular geometry, *perceptual epipolar disparity* is perceived.

These perceptual variations suggest that *geometry-based* and *context-based* depth cues contribute jointly rather than independently to depth representation.

To account for this joint contribution and to maintain consistent figure–ground organization across both physical and illusory domains, we extend the *layered disparity representation* introduced in Chen et al., (2025), which explicitly models the hierarchical integration of multiple interacting disparity sources.

#### Layered Structure of Disparity Representation

Disparity is not a uniform signal but a structured ensemble encompassing physical (epipolar), perceptual, and illusory (non-epipolar) components.

As established in Chen et al., (2025), the *layered disparity representation* provides a unified and flexible formulation that integrates absolute (Far/Near), motion-derived (optic-flow), and other disparity types within a single computational framework.

Conceptually, this representation accommodates both epipolar (physical) and non-epipolar (illusory) disparity formation, enabling coherent depth representation across spatial scales and disparity types. It also supports the propagation of border-ownership signals along both near- and far-surface boundaries. By maintaining consistent disparity sign and direction across layers, the model enables *hierarchical depth integration* and *stable ownership propagation* along object borders.

Computationally, *TcRd(k1*.*5)* realizes this principle through thresholded differencing of relative disparities in each layer, followed by cross-layer summation (Eq. 4.7). This produces border-ownership maps that remain consistent across physical and perceptual depth conditions.

Figure 9 illustrates this layered disparity representation, demonstrating how border ownership emerges from the combination of Far/Near disparities.

Finally, this *layered framework provides a natural substrate for higher-level modulation and feedback*, particularly from area V4, which will be further examined in Section 6.3.4 (Global Context Awareness), 6.3.7 (Thresholding) and 6.4.1 (V4 → V1 Feedback), where feedback-driven contour enhancement and cross-layer disparity interactions are discussed.

#### 6.3.2 Local Kernel and Long-Range Relative Disparity

Beyond the performance difference between the *k1*.*5* and *k3*.*4* kernels (Section 5.5.2(a)), their structural characteristics also diverge in how they encode spatial differencing.

The *k1*.*5* kernel satisfies a strong *additivity property*: the relative disparity between any two (horizonal) distant points can be obtained by summing the immediate-neighbor relative disparities along the connecting path (Figure 11). Formally,

**Figure 11.**
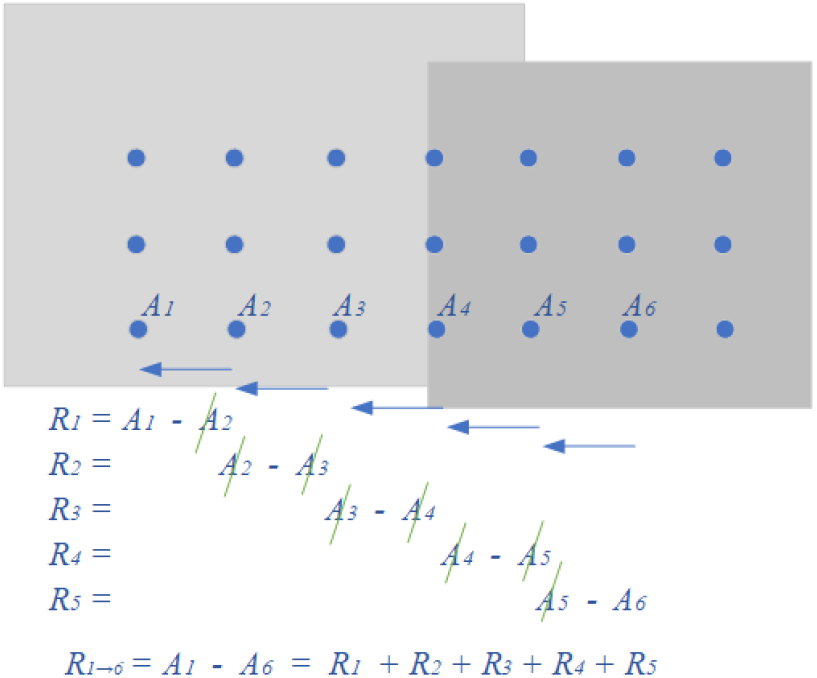
Relative Disparity between two distant points. The relative disparity between two spatially separated points can be computed by summing (or integrating) the immediate-neighbor relative disparities along the path connecting them. This illustrates the additive property expressed in Eq. (6.1)–(6.5). Here, *A*_*i*_ and *R*_*i*_ denote absolute and relative disparities at pixel *i*, respectively.

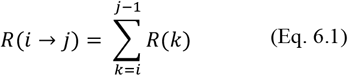

which implies a *transfer-like transitivity*,

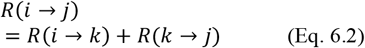

This ensures that *local disparity differencing scales coherently to long-range disparity relations*, enabling hierarchical depth integration across space.

When two distant points differ in both row and column directions, their total relative disparity can be decomposed into orthogonal components along the two orientations of the connecting path (row *r* and column *c*):

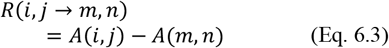

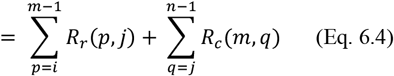

Here, *A*(*i, j*)and *A*(*m, n*)represent the layered disparities at the two points (*i, j*)and (*m, n*), while *R*_*r*_ and *R*_*c*_ denote the directed local relative disparities along rows and columns. This formulation expresses the total disparity difference between two locations as the accumulated sum of local disparity differences along any connecting path.

Because relative disparity under k1.5 obeys strict additivity, the computed *long-range disparity is path-independent*:

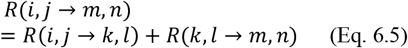

Path independence yields geometric consistency and stable depth relations across distances, enabling coherent long-range RD → BO correspondence from purely local differencing.

In contrast, the *k3*.*4* kernel lacks this additive consistency, resulting in partial loss of long-range coherence and degraded alignment between relative-disparity and border-ownership maps.

Although no direct neurophysiological evidence has yet established such additive behavior, related findings in V2—such as local pooling of multiple V1 disparity signals (Bredfeldt et al., 2006), context-dependent disparity tuning (Thomas et al., 2002), and hierarchical disparity integration over space and time (Kaestner et al., 2022)—suggest that *mechanisms with similar emergent properties are plausible*.

Both k1.5 and k3.4 are *linear difference operators* and therefore exhibit the Thomas-style equal-magnitude, same-direction shift (Thomas et al., 2002) when a comparison region’s disparity is uniformly offset. However, only k1.5 satisfies strict *additivity and path independence*, yielding more coherent correspondence between relative disparity and border ownership over long-ranges.

#### 6.3.3 Local Kernel and Owner-Side Direction

Directed relative disparity provides a natural biological basis for the owner-side direction, typically orthogonal to the object border (see Figure 3). However, detailed analysis indicates that this orthogonality is not always strictly maintained, whereas *side preference* remains consistent at the level of individual kernel components.

In general, the owner-side direction at an object-border pixel can be determined from the row- and column-wise components of its directed disparity difference, as shown in Figure 12 and (Eq. 6.6).

**Figure 12.**
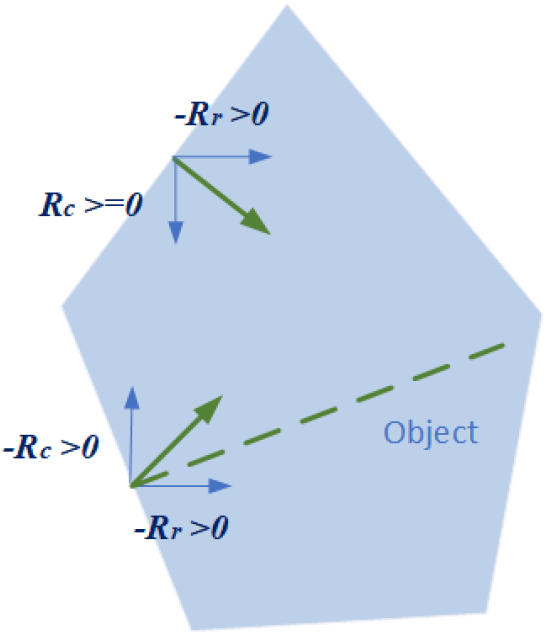
Owner-Side Direction from Local Kernel. In general, the owner-side direction (green arrows) on an object border can be determined from its row and column directed disparities (blue arrows), as shown. Notably, one owner-side vector deviates from the ideal orthogonal direction (green dashed line), illustrating that local variations in disparity structure can lead to non-perfect orthogonality while preserving consistent *side preference*.

Formally, the local owner-side direction vector 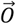 can be expressed as:

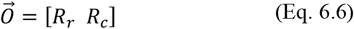

where *R*_*r*_ and *R*_*c*_ represent the row- and column-wise directed disparity differences at the border pixel. The vector 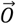 is consistent with the border-ownership vector representation defined in (Eq. 4.7).

| Component | Owner-side direction |
| --- | --- |
| $R_r \geq 0$ | left |
| $-R_r > 0$ | right |
| $R_c \geq 0$ | below |
| $-R_c > 0$ | above |

The sign of *R*_*r*_ or *R*_*c*_, determines the owner side—that is, the ***owner-side direction corresponds to the direction of increasing disparity***, indicating the region perceived as *nearer (the figure side)*.

*Deviations from perfect orthogonality are likely functional rather than accidental* (Figure 12), arising when disparity gradients are uneven, curved, or vary with local context—reflecting fine-scale variations in how neighboring neurons respond to visual inputs in area V2 (Zhou et al., 2000; Williford & von der Heydt, 2013).

In our previously proposed active-neuron–based surface-filling-in model (Chen et al., 2025), the *directional components* of the owner-side vector—its row and column elements—are *more critical* than the exact composite direction itself. These components guide the propagation of border-ownership signals along neighboring contour elements, ensuring consistent ownership continuity across a surface.

*This makes strict orthogonality of the owner-side direction even less critical*, as the underlying mechanism relies on local directional consistency rather than geometric precision.

Although direct physiological evidence is not yet available, it remains plausible that neurons with distinct structural configurations could implement the required directional components through specialized subfields or other arrangements. Such mechanisms would allow flexible handling of owner-side direction and orthogonality. The precise neural mechanisms, however, remain to be clarified and deserve further investigation.

#### 6.3.4 Local Kernel and Global Context Awareness

Local disparity differencing (via kernels such as k1.5) governs the fine-scale computation of relative disparity and border-ownership, but these operations do not operate in isolation.

In our prior work (Chen et al., 2022; Chen et al., 2025), we proposed that border ownership and category selectivity both require *global context awareness*— contextual guidance that extends beyond the classical receptive field to achieve coherent global percepts.

We further suggested that this context awareness is embedded within border-ownership– and category-selective neurons in V2, enabling them to integrate local disparity cues with broader contextual signals while still matching the short latency observed of border-ownership responses (∼ 25 ms; Zhou et al., 2000).

The view was supported by TcNet experiments in which the network learned and applied global context, effectively emulating “inherently knowledgeable” neurons that behave like a self-sufficient perceptual ‘mold’.

The present study extends this framework by showing that *border ownership emerges as thresholded relative disparity*, which naturally implies that the ***computation of relative disparity itself is subject to global contextual modulation***.

This relationship is empirically supported by the comparison experiments (Section 5.5.2(b); Table 2, Rows 19 ≈ 21 ≈ 95% for border ownership), demonstrating that *TcNet* trained on disparity inputs achieves performance comparable to *TcRd(k1*.*5)*—a training-free model that directly implements RD sign coding (Figure 4(b)).

These findings indicate that local disparity differencing and global contextual modulation act cooperatively, with the latter guiding the consistency and stability of relative-disparity-based border-ownership representations.

#### Modulation by Global Context Awareness

Further analysis suggests that the contextual modulation of relative disparity may occur through two complementary pathways:

##### Indirect modulation via V1

Global contextual influences may shape the *layered disparity representations* formed in V1, thereby producing the structured disparity patterns that V2 subsequently processes through differencing and thresholding to generate the expected border-ownership signals.

##### Direct modulation within V2

Alternatively, or additionally, global context awareness embedded in V2 itself may directly guide border-ownership generation by integrating contextual priors with local disparity inputs.

Together, these mechanisms provide a functional bridge between local kernel operations and global perceptual coherence, forming the foundation for the feedback-driven contextual modulation further analyzed in Section 6.4.

#### 6.3.5 Category Selectivity

In our prior work (Chen et al., 2022; Chen et al., 2025), we proposed that category selectivity at early visual stage is jointly generated with border ownership under the guidance of global context awareness.

This view aligns with neuroscientific findings by Silson et al. (2022), who demonstrated that the human visual cortex is organized broadly according to two major principles—***retinotopy and category selectivity***—and that *retinotopic maps and category-selective regions show considerable overlap*.

If border ownership indeed arises from thresholded relative-disparity computation, then the co-localized ***category selectivity*** observed by Silson and colleagues (2022) is plausibly generated through a ***similar mechanism***.

In this interpretation, the *category selective channel* described in our prior work (Chen et al., 2022; Chen et al., 2025) provides an efficient means by which global context awareness identifies and reinforces borders belonging to a given category, while the co-located border-ownership channels encode their spatial positional and relative depth.

This functional coupling naturally explains why *category selectivity and border-ownership generation are co-localized* in early visual areas, *reflecting a shared computational substrate* for contextual segmentation and depth organization, thereby *avoiding computational redundancy*.

#### Category Illusory Disparity

From a neuroscientific perspective, a plausible and biologically motivated hypothesis is that *category-associated, disparity-like “illusory object blocks”—*here referred to as *Category Illusory Disparity—*may be integrated into the layered disparity representation in V1 through global contextual modulation.

These structured, context-driven disparity layers could then undergo *disparity differencing in V2*, producing category-specific contours within the same computational framework that governs depth and ownership processing.

Although direct neurophysiological evidence for this mechanism is not yet available, it is partially supported by our computational simulations, which exhibit analogous behavior in the model.

Additional support comes from recent fMRI work showing that category rules can dynamically reshape population representations even in early visual cortex (Henderson et al., 2025), suggesting that contextual or task-dependent categorical structure can be embedded at retinotopic stages prior to higher-level object recognition.

Together, these observations offer a coherent theoretical account of how category information might be embedded within disparity-based computations of figure– ground organization.

(See ‘Illusory Disparity’ in Section 6.3.1 and the discussion of V4-mediated feedback in Section 6.4)

### Contextual vs Content Categorization

This perspective suggests a natural division:

- *Contextual categorization*—an early, spatially grounded category assignment driven by contextual priors and integrated with disparity-based segmentation
- *Content categorization*—a later, higher-level category processing that abstracts object identity independent of precise spatial layout.

This joint configuration supports rapid, contextually informed segmentation, providing a biologically plausible foundation for the **B + C** (border-ownership + category) representation used in the present framework.

#### 6.3.6 Relative-Disparity and Border-Ownership Coding Configuration

Border-ownership (BO) generation can be implemented through distinct coding configurations that differ in how directional selectivity and ownership side are represented.

Two main configurations are employed in this study— *opposite-channel (OC)* coding and *relative-disparity (RD)*-based coding.

##### RD-based vs OC-based Coding

If the hypothesis that *thresholded relative disparity represents the neural basis of border ownership* is correct, then the RD-based configuration provides the more biologically plausible interpretation.

This view is supported by comparison experiments using both RD-based and OC-based ground-truth variants of the VKitti dataset (see Section 5.5.2(b); Table 2, Row 19 ≫ Row 11 and Row 20 ≫ Row 15), where RD-based coding exhibited superior correspondence specifically when the disparity–ownership structure was internally consistent, confirming that RD-based coding aligns more naturally with the underlying depth structure.

In contrast, RD-based and OC-based codings are effectively identical for the RDS dataset, whose depth structure follows symmetric epipolar geometry.

From a computational perspective, given physical, illusory, or even pseudo-disparity ground truth, the RD-based configuration is simpler and more robust, as RD-based border ownership ground truth can be obtained via kernel-based disparity differencing, whereas OC-based coding requires explicit and often complex contour separation and ownership-side assignment.

This simplicity, combined with its biological plausibility, makes RD-based coding a unified and efficient representation for border-ownership generation across both physical and illusory depth domains.

Finally, because of the close correspondence between OC coding and RD sign coding (Figure 4(b)), the raw pre-threshold outputs of TcNet models trained on OC-based ground truth can be approximately interpreted as relative-disparity maps, following RD sign configuration.

In this sense, the network implicitly learns the relative-disparity structure even when trained on OC labels.

##### Among RD-based Codings

As described in Section 5.1, the 4-channel and 2-channel orientation RD coding (Figure 4(a), (c)) represent distinct implementations of relative-disparity differencing and thresholding without overlap between row and column components, resulting in clean and non-redundant representations.

The 4-channel configuration (Figure 4(a)) explicitly separates horizontal and vertical disparity differences for both positive and negative owner-side directions, providing full directional selectivity at the cost of increased computational and representational resources.

By contrast, the 2-channel orientation configuration (Figure 4(c)) encodes only signed disparity differences along the principal axes, effectively collapsing directionally opposite responses into paired ownership channels.

This simplification preserves ownership-side information while reducing representational redundancy, making the *2-channel RD orientation coding* both computationally efficient and biologically plausible.

Although RD-based coding provides a coherent computational interpretation of border-ownership signals, the precise neuronal or microcircuit mechanisms by which biological vision implements such coding remain unknown. Determining how neurons encode relative disparity and ownership side is an open question that warrants further investigation.

#### 6.3.7 Thresholding and Border Contour

In our experiments, the threshold values for border ownership were empirically selected to be balanced— sufficiently high to suppress noise yet low enough to retain valid contour detections. However, regardless of the threshold value, the resulting output may not always form thin, single-pixel contours. To enable fair benchmark comparison, a post-processing “thinning” step was applied to the border-ownership maps (see Section 5.4).

This raises an important computational and biological question: ***must border ownership representations in V2 form thin borders?***

From a computational perspective, thresholding over relative disparity, as illustrated in Figure 9, can produce continuous regional activations (e.g., large patches of distant ground or foliage) whose disparities exceed threshold. Such region-like responses, though not geometrically thin, can still effectively guide ownership propagation within the surface-filling-in model proposed in our prior work (Chen et al., 2025), albeit with potentially reduced efficiency.

Although no neurophysiological study has yet established whether border-ownership representations in V2 consistently form thin, contour-like patterns, such an outcome—if eventually verified—would imply the involvement of an additional disparity-processing mechanism beyond simple local thresholding.

Such a mechanism could *selectively suppress broad regional activations while enhancing contour transitions*. This behavior could emerge from intrinsic feedforward computations, but feedback from higher-level area (e.g., V4) provides a compelling and biologically plausible modulation pathway. The layered disparity representation discussed earlier (Section 6.3.1) offers an ideal substrate for such feedback integration, enabling V4-driven enhancement or suppression across disparity layers to refine border precision and contextual consistency.

This possible feedback modulation will be further examined in Section 6.4 (Roles of V4 in Feedback).

#### 6.3.8 Feature Foundation for Layered Disparity and Border Ownership

These computations presuppose an earlier stage of feature extraction—such as edge, orientation, contrast, color, and texture processing (Hubel & Wiesel, 1959; Carandini et al., 2012; Livingstone & Hubel, 1984; Freeman et al., 2013)—but it remains unclear to what extent these features directly contribute to the formation of absolute-disparity, motion parallax signals, or to the subsequent relative-disparity and border-ownership computations.

Within the present framework, ***disparity-based mechanisms appear to dominate RD and BO generation***, whereas feature-derived signals are likely to play a more indirect or downstream role, most plausibly in surface-related processes such as surface filling-in (Chen et al., 2025), rather than in the core depth-based segmentation mechanism itself.

A useful empirical indication comes from Section 5.5.2(e) (Disparity with and without Stacked RGB).

While exploratory in scope, these experiments nevertheless provide insight into how feature cues may interact with disparity-based computations.

For the **RDS** dataset, strong in disparity but weak in contextual structure, TcNet models trained with *stacked RGB plus disparity* inputs outperformed models using disparity alone.

This suggests that when stimuli are context-poor or contain minimal scene structure, additional feature cues (e.g., local luminance/color contrast) can help stabilize disparity estimation or enhance border localization, thereby improving the downstream RD and BO computations.

In contrast, for the photorealistic **VKitti** dataset, models trained with and without stacked RGB achieved comparable performance.

This indicates that, once high-quality Far/Near absolute disparities have been derived from the RGB images, these disparity maps already capture the essential depth information required for RD computation and BO generation. In other words, for naturalistic scenes where disparity extraction is reliable, adding RGB at the TcNet input stage does not substantially change RD or BO performance.

Taken together, these findings support the following picture:

- *Layered disparity channels are the primary substrate* for RD and BO generation.
- *Feature cues (RGB / edges / texture)* help mainly when disparity signals or contextual structure are weak (as in RDS), by supporting more robust disparity estimation or edge localization *upstream* of RD and BO computation.
- Once layered disparities are well-formed, *relative disparity differencing and thresholding dominate border-ownership generation*, with features contributing only indirectly via their role in disparity and motion estimation.

Thus, while feature extraction in V1 (e.g., orientation, color, contrast) is a necessary foundation for building layered disparities and optic-flow signals, our results indicate that ***the core mechanism for border ownership and depth-based figure–ground organization operates at the level of disparity***, rather than at the level of features per se.

Although the precise contribution of specific feature channels to disparity estimation, BO generation and surface filling-in (Chen et al., 2025) remains an *open question*, our findings support the conclusion that layered disparities, rather than raw feature maps, constitute the primary substrate for the RD- and BO-centered computations analyzed in Sections 6.3 (Nature and Structure of Disparity) and 6.4 (Roles of V4 in Feedback).

### 6.4 Roles of V4 in Feedback

Although the feedforward computation of thresholded relative disparity in V2 provides a fast and powerful basis for border ownership and depth-based figure–ground organization, several phenomena indicate that *feedforward mechanisms alone are insufficient to account for full perceptual coherence*, suggesting a role for additional modulation from higher visual areas.

First, *local thresholding of relative disparity can produce region-level activations*, especially in smooth depth-transition regions or extended depth fields where relative-disparity differences remain above threshold across a broad spatial area (Section 6.3.7). While such responses still support ownership propagation (Chen et al., 2025), it remains unknown whether biological vision naturally produces similarly broad, region-like border-ownership signals—a question that has not yet been resolved in neurophysiology.

If future evidence were to show contour-focused border-ownership selectivity, this would imply the presence of *top-down mechanisms that refine, suppress or enhance border precision* in ways that complement the feedforward differencing.

Second, *illusory, perceptual, or non-epipolar disparity components* (Sections 6.3.1 and 6.3.5) are strongly influenced by global scene interpretation and contextual priors. Although such disparities may originate partially through local computations, their stability, strength, and perceptual dominance appear to depend on broader *contextual reinforcement* beyond the capacity of V2’s local mechanism alone.

Together, these considerations motivate a ***hierarchical recurrent architecture*** in which *V4 supplies contextual, categorical, and precision-enhancing signals to V1 and V2*. This feedback refines layered disparities, stabilizes ambiguous or multistable borders, and aligns early depth computation with global perceptual organization.

Consequently, *V4 plays a crucial role in shaping the dynamic, context-aware figure–ground representations* examined in the following subsections.

#### 6.4.1 V4 → V1 Feedback and Layered Disparities

##### Structured Feedback and Functional Role

Recent evidence suggests that V4 provides structured feedback to V1 that modulates early-stage feature processing in a manner aligned with figure–ground organization and border-ownership tuning (Jeurissen et al., 2024).

In that study, neurons in V4 showing figure–ground selectivity (functionally analogous to border-ownership selectivity) were shown to establish *targeted feedback projections* to V1 domains with matching orientation tuning and figure-ground preference, effectively reinforcing figure-side contrast while suppressing background regions.

##### Feedback Acting on Layered Disparities

This top-down feedback provides a *biologically plausible mechanism* for modulating the *layered disparity representation* used in our framework (Section 6.3.1; and Chen et al., 2025). Because the layered disparity representation in V1 serves as the input from which V2 computes relative disparity and border ownership, ***any modulation of these layers by V4 directly influences the subsequent RD and BO signals*** generated in V2.

V4 feedback could modulate individual disparity layers, enhancing contours at strong depth discontinuities and attenuating redundant or smooth disparity fields.

This would yield more contour-focused border-ownership activations, consistent with enhanced figure-ground modulation observed in V2 under figure–ground conditions (Fang et al., 2009; Jeurissen et al., 2024).

##### Category-Consistent Modulation

Although direct neurophysiological evidence is lacking, it is plausible that category selectivity interacts with the layered disparity representation through the same feedback mechanisms that refine border ownership and depth organization.

In this view, V4 → V1 feedback may reinforce *category-consistent disparity layers*, instantiating or biasing the re-weighting of layered disparities toward contours associated with category-relevant structures while attenuating incongruent ones.

Such modulation would link early disparity refinement with higher-level categorical priors, providing a plausible pathway through which category-induced illusory disparities (Section 6.3.5) become integrated into the layered disparity framework.

This feedback-driven interaction supports an efficient mechanism for aligning early figure–ground computation with perceptual categorization (see also Section 6.4.2 for contextual stabilization).

##### Perceptual Alternations: Explanation

Importantly, *perceptual alternations* such as those illustrated earlier in Figure 10 and the Rubin Face–Vase illusion (Figure 13) can be interpreted as *manifestations of the dynamic role of this feedback loop*.

**Figure 13.**
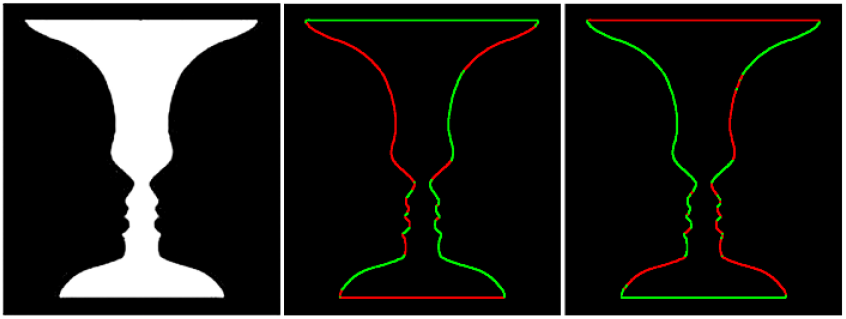
Rubin Face-Vase Illusion (adapted from Fig. 11 in Chen et al., 2025). **Left**: Rubin Face-Vase image (source: Google); **Middle**: Two-channel OC-based border-ownership map when perceived as two faces. **Right**: Two-channel border-ownership map when perceived as a vase.

The same 2-D input can support multiple perceptual organizations—illusory, perceptual-epipolar, or physical depth interpretations—depending on how disparity layers are re-weighted, instantiated or combined over recurrent cycles.

Such perceptual switching suggests that higher-level areas like V4 can trigger *reorganization of disparity representations in V1*, effectively modulating border-ownership structures and figure–ground relationships in semi-real time.

Similarly, the escalated depth in Kanizsa illusions (Figure 7) may plausibly arise from *V4-mediated amplification and integration* of illusory (non-epipolar) disparity layers feeding back into V1, reinforcing local contrast and border signals to yield heightened depth salience.

##### Perceptual Alternations: Natural Plausibility and Theoretical Novelty

Unlike previous interpretations that attribute perceptual alternations primarily to higher-order attention, competitive cortical dynamics, or value-driven perceptual decision processes (Logothetis et al., 1996; Tong et al., 1998; Leopold et al., 1999; Safavi et al., 2022), the present framework explains them as a ***natural consequence* of *disparity-layer selection and recurrent re-weighting*** within the established V4 → V1 feedback loop.

Because the layered disparity representation is already instantiated in early visual stages, selective feedback only needs to re-weight and combine existing disparity layers rather than regenerate entirely new percepts, making alternations both *biologically economical* and *temporally plausible*.

This interpretation aligns with short perceptual cycle times and known cortical latency patterns, suggesting that bistable percepts such as the Rubin Face–Vase and Kanizsa illusions arise from *feedback-driven re-weighting and integration* among existing depth layers, providing a ***unified and mechanistically grounded explanation for figure–ground multistability***.

##### Feedforward and Delayed Feedback

Conceptually, this mechanism operates much like a ***slideshow***: each disparity layer represents a “slide,” depicting a distinct perceptual organization—illusory, perceptual-epipolar, or physical (with physical layers serving as the default representation).

Feedback from V4 acts as the *selector and modifier*, bringing one or multiple slides into prominence while fading others, orchestrating which depth configuration dominates the *next perceptual cycle*.

Because border-ownership generation in V2 occurs with short latency (∼25 ms; Zhou et al., 2000; Chen et al., 2025), V4 feedback likely influences *subsequent rather than concurrent* processing stages, refining and reinforcing the perceptual outcome in the *next iteration*.

In this view, perception evolves as a *continuously updated sequence of selections and combinations*—each cycle shaped by *both* ***feedforward thresholded disparity differencing*** *and* ***delayed feedback modulation***.

##### Higher-Level Modulatory Control

Although the exact source of top-down modulation is beyond the present paper’s scope, *V4 feedback can be regarded as a principal hypothesized pathway* through which higher-level visual or contextual control influences early visual processing.

Acting through its feedback projections, V4 dynamically re-weights disparity layers and border-ownership signals in V1 and V2—enhancing contours associated with the dominant perceptual interpretation while suppressing weaker or redundant activations. In this sense, the *feedback acts as a selective control mechanism* that determines which disparity configuration is reinforced in subsequent processing cycles, without altering the intrinsic short-latency feedforward computation of border ownership.

##### Hierarchical Recurrent Loop

**V4 → V1 feedback** therefore functions *not merely as a refinement process but as a dynamic selection mechanism*, capable of toggling and combining different disparity layers to determine the dominant percept. It serves as a *precision-control and perceptual-selection mechanism* superimposed on the feedforward thresholded disparity differencing implemented by local operators such as k1.5:

*V1(layered disparities)* → *V2 (local differencing and thresholding)* → *V4 (ownership and percept selection)* → *V1 (feedback reinforcement of selected layers)* (see also Figure 14)

**Figure 14.**
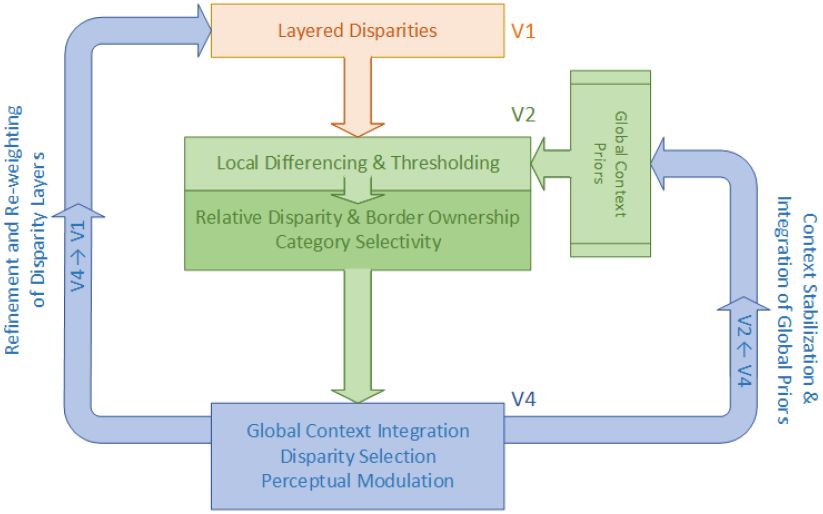
V4 Feedback Flow Diagram. Schematic illustration of dual V4 feedback pathways. The **V4→V1 feedback** primarily *refines and re-weights* layered disparity representations, enhancing contour precision and depth coherence. In contrast, the **V4→V2 feedback** provides *contextual stabilization and integration*, incorporating global context priors such as object continuity and category consistency to align local border-ownership and category-selective signals with the dominant global interpretation. A broader overview of the unified architecture is shown in Figure 15.

This *hierarchical recurrent loop* explains how feedback can both sharpen contour precision and enable perceptual reversals—as seen in bistable or illusory configurations—thereby *uniting the dynamic aspects of disparity, border ownership, and perceptual awareness*.

Collectively, this framework links feedback-based figure–ground modulation with the layered disparity architecture proposed here and in Chen et al. (2025), providing a mechanistic account of perceptual reversals such as the Rubin Face–Vase and Kanizsa illusions within a unified recurrent model.

#### 6.4.2 V4 → V2 Feedback and Global Context Awareness

Whereas V4 → V1 feedback governs the temporal refinement and selection of disparity layers across perceptual cycles, its complementary pathway—V4 → V2—serves a distinct role in spatial and contextual integration within the figure-ground hierarchy.

##### Global Context Awareness

In our prior work (Chen et al., 2022; Chen et al., 2025), we introduced the concept of *global context awareness or prior knowledge*, referring to the visual system’s capability to incorporate contextual information across wide areas of the visual field when computing border ownership.

Because border-ownership signals emerge with extremely short latency (∼25 ms; Zhou et al., 2000), we proposed that this capacity is best understood as an intrinsic property of border-ownership– and category-selective neurons rather than the outcome of slower recurrent inference.

These neurons, *distributed* across V2, collectively encode figure–ground organization in a *context-sensitive yet rapid* manner, enabling globally coherent ownership assignments even under ambiguous or multistable conditions.

Over time, this *built-in contextual sensitivity* could *refine* through exposure to new visual experiences or environmental changes, allowing the system to generalize figure–ground organization across objects, categories, and viewpoints.

##### Anatomical Basis for V4 → V2 feedback

Anatomical evidence strongly supports *dense feedback connections from V4 to V2*, consistent with a hierarchical recurrent architecture.

Large-scale cortical tracing in macaque visual cortex (Markov et al., 2014; Franken et al., 2021) demonstrated that *V4 provides one of the dominant feedback inputs to V2*, projecting primarily from deep cortical layers (L5–L6) and terminating in the superficial and deep layers of V2— the laminar distribution characteristic of feedback modulation.

These projections provide the *structural substrate* through which V4 can deliver *context-based modulation* signals that refine local disparity differencing and ownership selectivity in V2 after the initial feedforward sweep.

##### Functional Implications

Functionally, V4→V2 feedback enables the integration of *global contextual priors*—such as shape continuity, object identity, and category consistency—into *local* ownership computations.

The V4→V2 feedback and the associated global contextual priors thus jointly act as a *contextual stabilizer*, with *feedback updating the priors* and the *priors biasing* ownership assignments toward perceptually dominant interpretation and *aligning* local disparity-derived ownership signals with the globally consistent interpretation selected at higher cortical levels (see also Section 6.4.1).

Through this mechanism, the visual system maintains *perceptual stability* even during ambiguous or bistable scenes, ensuring that border ownership and figure–ground organization converge toward a *coherent, context-consistent percept* across successive processing cycles.

##### Hierarchical Functional Loop

*V1 (layered disparities)* → *V2 (local border-ownership and category selectivity generation)* ↔ *V4 (global context integration and feedback modulation)* (see also Figure 14)

Together with V4 → V1 feedback, this pathway completes a *bidirectional recurrent loop* that balances fine-scale disparity precision with large-scale contextual coherence—forming the functional backbone of dynamic border-ownership computation in the primate visual hierarchy.

#### 6.4.3 Dual Feedback Roles of V4

The analyses above suggest that *V4 serves as a dual-feedback hub*, providing complementary modulatory influences to both early visual areas—*V1 and V2*—within a recurrent cortical hierarchy.

**V4 → V1 Feedback** primarily refines the *precision of layered disparity representations*, sharpening contour selectivity and enabling perceptual switching and combination among different depth configurations (illusory, perceptual-epipolar, or physical).

Through this pathway, V4 enhances the fidelity of depth and border cues at their earliest representational stages, converting diffuse disparity activations into enhanced, contour-like border-ownership maps (see Section 6.4.1).

**V4 → V2 Feedback**, in contrast, provides *contextual stabilization and refinement*, integrating global context priors such as object continuity, category consistency, and scene-level coherence.

This feedback aligns local ownership assignments with the dominant global interpretation selected at higher cortical levels, ensuring perceptual consistency and figure–ground coherence across successive processing cycles (see Section 6.4.2).

Together, these reciprocal pathways implement a *hierarchical recurrent loop* that balances *precision* and *context* (also see Figure 14):

*V1 (layered disparities)* → *V2 (local relative disparity & border-ownership generation and category selectivity)* ↔ *V4 (global context integration and selective feedback)* ↔ *V1 (refinement and re-weighting of disparity layers)*

This integrated framework positions **V4** as a *dynamic control node* in the border-ownership network— *stabilizing perceptual context through feedback to V2 while simultaneously refining disparity and border ownership precision through feedback to V1*.

The resulting architecture unites the *feedforward and feedback* components of figure–ground computation into a coherent, biologically plausible model of *adaptive perceptual organization*.

### 6.5. A Unified Depth Perception and Figure-Ground Organization Model

Building on the expanded theoretical and empirical findings presented in this study, we refine and extend the *Border Ownership Centered Figure–Ground Organization Model* originally proposed in Chen et al. (2025).

The updated model (Figure 15) integrates the layered disparity representation, thresholded relative-disparity differencing, and dual V4 feedback pathways and active-neuron dynamics into ***a coherent, biologically grounded framework* for *depth perception* and *figure–ground organization***.

**Figure 15.**
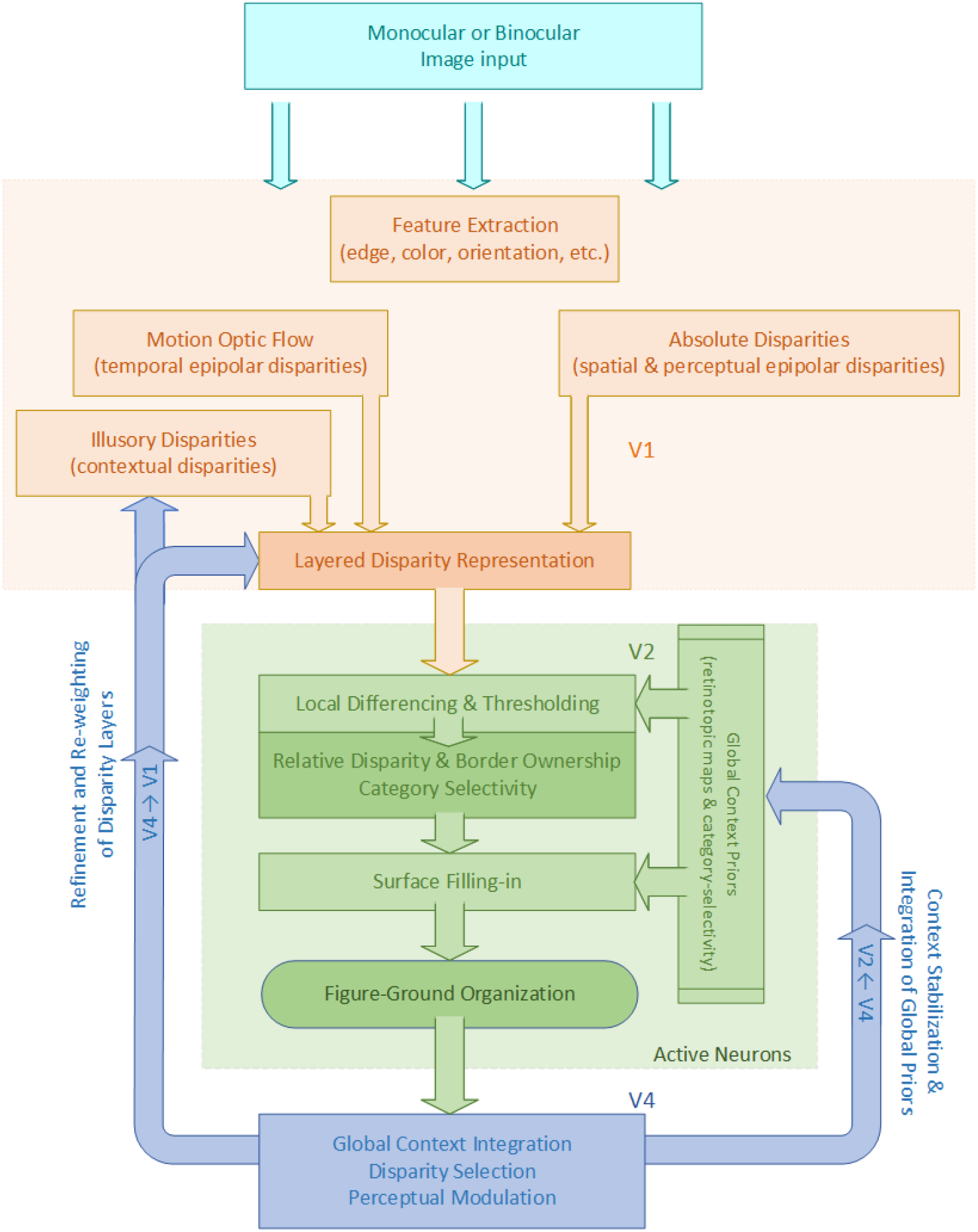
Architecture Diagram of a Unified Depth Perception and Figure-Ground Organization Model. Schematic illustration of the unified framework proposed in this study. The model integrates: (i) a **feedforward mechanism** in which **border-ownership and early category selectivity** arise intrinsically from thresholded relative-disparity differencing; (ii) a **layered disparity representation** that accommodates physical (epipolar), perceptual-epipolar, and illusory/contextual disparity components; (iii) **dual V4 feedback pathways** (**V4→V1 and V4→V2)** that jointly coordinate precision and contextual organization; (iv) active-neuron-mediated **surface filling-in** for completion of figure-ground organization; and (v) low-level feature extraction (edge, color, orientation, etc.) depicted as a V1 substrate for completeness—its direct contribution to layered disparities, RD, and BO generation remains uncertain (Section 6.3.8).

#### 6.5.1 Feedforward Framework: Intrinsic Relative Disparity, Border-Ownership and Early Category Selectivity

At the core of the model is the central finding of this work: *border ownership arises intrinsically through thresholded relative-disparity differencing* (Section 6.3).

This same feedforward mechanism also provides the structural foundation for *early category selectivity*, consistent with our prior proposal that category signals and border-ownership signals share a common contextual basis and emerge in co-localized circuits (Section 6.3.5).

This feedforward mechanism:

- *operates concurrently with border formation*, because the border itself emerges from a thresholded disparity difference;
- *applies uniformly across* contrast-defined, disparity-defined, and illusory/contextual borders;
- *encodes* both *directional selectivity* and *ownership polarity* at each border pixel; and
- *provides* the stable depth-anchored substrate over which category-selective biases can be imposed, allowing contextual or category-consistent signals to align naturally with disparity-structured borders.

In this view, the feedforward computation provides an *early, self-sufficient* transformation from disparity structure into directed ownership and early categorical bias within *short latency*.

This forms the foundation for the later *surface-filling-in* processes and for the *global, recurrent figure–ground organization* refined through V4-mediated feedback (Sections 6.5.2-6.5.4).

#### 6.5.2 Layered Disparity Representation: The Structural Core

A major refinement over the *layered disparity representation* introduced in our prior work (Chen et al., 2025) is its explicit expansion to accommodate physical (epipolar), perceptual-epipolar, and illusory (non-epipolar) components within a single depth-encoding framework (Section 6.3.1).

This structure allows depth cues that are *geometric, perceptual, or context-driven* to be placed into a unified representational system, enabling the model to handle real, ambiguous, and illusory depth on *equal computational terms*.

Within this structure, feedforward relative-disparity differencing is computed jointly over the entire layered representation, generating a multi-layer border-ownership code in a single operation. The resulting border-ownership signals are organized by layer, while cross-layer integration:

- maintains *global depth coherence*,
- supports *transitions* between 2-D and 3-D interpretations, and
- enables *perceptual alternations* such as the Rubin Face–Vase depth reversals.

Because of this organization, the layered representation provides a *natural substrate for hierarchical feedback*, particularly from V4, which can selectively enhance, suppress, or re-weight depth layers according to perceptual dominance or task demands.

#### 6.5.3 Recurrent Feedback Integration: Precision and Context Modulation

Building upon the dual feedback mechanisms analyzed in Section 6.4, the updated model incorporates two *complementary feedback loops* that jointly regulate disparity structure, border ownership, and global perceptual organization.

**V4 → V1 Feedback** (Precision and Layer Selection): This pathway operates primarily on the *layered disparity representation*, where it:

- *refines and re-weights* the layered disparity maps,
- *sharpens* contour precision at depth discontinuities,
- *attenuates* homogeneous or redundant disparity fields, and
- enables *switching and recombination among multiple disparity layers to support flexible perceptual organization*.

Critically, this modulation can act not only by *re-weighting* existing layers but also, over successive processing cycles, by *instantiating or strengthening illusory or context-driven disparity* components (e.g., Kanizsa-like illusory or category illusory disparity).

Under this view, perceptual reversals and depth “pop out” effects arise from *dynamic reconfiguration of a disparity ensemble*—sometimes by shifting weights among layers, sometimes through the emergence of new contextual or illusory layers.

This mechanism thus provides a natural and biologically plausible account of bistable and multistable percepts (e.g., Rubin Face–Vase), in which competing figure–ground organizations correspond to different combinations of disparity layers selected and reinforced across recurrent cycles.

**V4 → V2 Feedback (**Context Stabilization and Evolution): This pathway provides *global contextual stabilization*, integrating *higher-level priors* (e.g., Gestalt principles) such as:

- shape continuity and object coherence,
- category consistency, and
- scene-level organization.

Through this integration, V4 feedback aligns local border-ownership signals with the *dominant global interpretation*, maintaining perceptual stability even when depth cues are ambiguous or conflicting. At the same time, this pathway enables perceptual organization to *evolve dynamically* as contextual information or task demands shift.

##### Precision-Context Integration

Together, the dual feedback pathways implement a *precision–context modulation architecture*:

- **V1** supplies high-fidelity, layered disparity structure (including physical, illusory, and category-related components),
- **V2** computes relative disparity, border ownership and category selectivity from this multi-layer representation, and
- **V4** modulates both according to global perceptual organization, category consistency, and scene-level priors, thereby governing the stability and evolution of the overall figure–ground interpretation.

#### 6.5.4 Active-Neuron Dynamics and Surface Filling-I

As proposed previously (Chen et al., 2025), border-ownership–selective neurons in V2 function as distributed “*active neurons*”, propagating ownership signals along the owner-side direction and maintaining object-centered continuity.

Under the unified model:

- The tight coupling between relative disparity and border ownership provides a more stable and error-resistant basis for ownership propagation,
- near- and far-surface continuity is preserved across depth layers, and
- seamless figure–ground organization persist during dynamic, ambiguous, or bistable viewing.

This mechanism forms the operational bridge between border-ownership generation and surface filling-in, ensuring coherent depth structure and surface continuity across perceptual cycles and feedback-driven updates.

Together with the layered disparity representation and dual V4 feedback pathways, the active-neuron mechanism completes *a hierarchical and biologically plausible cycle from depth encoding to stable figure– ground organization*.

## 7. Limitations and Future Directions

Although the present framework outlines a unified account of disparity, border ownership, and feedback interactions, several important questions remain.

**First**, although the proposal that border ownership arises from thresholded relative-disparity differencing provides a concise explanation for many classical BO properties (Section 6.2), the neural circuitry capable of implementing this computation is not yet established. Determining whether V2 connectivity can support such directed differencing will require targeted physiological and computational investigations.

**Second**, some components of the model—particularly the contributions of illusory and category-dependent disparity signals—remain theoretical and are currently supported only indirectly by computer-vision simulations. Whether early visual cortex explicitly represents such signals, or whether they instead arise through top-down modulation, remains to be determined.

**Third**, the functional roles assigned to V4 feedback— precision modulation in V1 and contextual stabilization in V2—remain incompletely characterized. Establishing the specificity, timing, and necessity of these pathways will require temporally resolved and laminar-specific experiments.

**Fourth**, the current formulation focuses primarily on static images and well-defined object boundaries. Natural scenes often contain complex, extended, or texture-rich contours—such as foliage, distant ground surfaces, or low-contrast boundaries—where border thickness, scale, and spatial uncertainty may affect ownership computation. How thresholded disparity differencing operates on such broad or distributed edges remains to be determined. Extending the framework to incorporate temporal dynamics, motion cues, adaptive receptive-field scaling, and eye-movement–dependent integration will be essential for evaluating its generality in ecological viewing conditions.

**Finally**, the proposed link between border ownership and surface filling-in, instantiated through active-neuron dynamics, is supported primarily by computational evidence. Direct tests of ownership propagation and its temporal evolution in cortex remain an important direction for future work.

Overall, the framework yields a set of clear, testable predictions regarding the relationship between disparity structure, ownership polarity, and feedback modulation. Evaluating these predictions in physiological and psychophysical experiments will be critical for determining the degree to which the proposed mechanisms reflect neural computation in natural vision.

## 8. Summary

This study proposes ***a unified framework for depth perception and figure–ground organization*** that integrates feedforward disparity computation, layered depth representation, border-ownership generation, and recurrent cortical feedback.

The *central insight* is that ***border ownership arises intrinsically from thresholded relative-disparity differencing***, yielding a short-latency feedforward transformation from local disparity structure to directional ownership. Recurrent feedback modulates and stabilizes these signals but is not required for their initial generation. This feedforward mechanism generalizes across contrast-defined, disparity-defined, and illusory borders, revealing a *common computation underlying diverse perceptual phenomena*.

Building on this foundation, we propose an expanded ***layered disparity representation*** that accommodates physical (epipolar), perceptual-epipolar, and illusory depth cues within a single coherent system. This representation serves as the structural backbone for both border-ownership coding and hierarchical feedback. Through cross-layer integration, it supports transitions between 2-D and 3-D interpretations, maintains global depth coherence, and provides a natural substrate for perceptual alternations such as the Rubin Face–Vase.

The framework further incorporates a ***dual V4 feedback architecture*** involving V4→V1 precision modulation and V4→V2 contextual stabilization. These pathways selectively enhance, re-weight, or suppress depth layers according to perceptual demands, shaping figure–ground organization over successive perceptual cycles while preserving the intrinsic feedforward computation.

Finally, the ***active-neuron dynamics*** in V2 (Chen et al., 2025) provide a mechanistic bridge between border ownership and surface filling-in: BO supplies the intrinsic owner-side direction that drives ownership propagation along object boundaries, while RD-supported disambiguation ensures coherent object-centered continuity.

Taken together, the proposed model provides a biologically plausible, computationally efficient, and conceptually unified account of depth perception and figure–ground organization. By linking disparity structure, border-ownership signals, feedback modulation, and perceptual multistability within a single framework, this work offers new insights into how the visual system constructs stable percepts from ambiguous and incomplete sensory inputs, and lays a foundation for future investigations into the neural and computational principles underlying perceptual organization.

## Author Contributions

- **Tianlong Chen**: Conceptualization, methodology, experimental design and conduction, data preparation, result analysis and discussion, writing, and editing.
- **Xuemei Cheng**: Data preparation, result discussion, paper review, and providing critical feedback.

## Data Availability

Both the Random Dot Stereograms dataset, the modified Virtual KITTI 2 dataset and the Kanizsa illusory contour dataset generated and analyzed during this study (Section 5) are publicly available on Zenodo under the following citations:

Chen T. (2025). *Border Ownership and Category Annotation Extended from Virtual KITTI 2* [Data set]. Zenodo. https://doi.org/10.5281/zenodo.15375453

Chen, T. (2025). *Random Dot Stereogram (rds4) Dataset with Border Ownership Annotation* (Version v1) [Data set]. Zenodo. https://doi.org/10.5281/zenodo.16423201

Chen, T., & Cheng, X. (2026). Kanizsa Illusory Contour (kani2) Dataset with Border-Ownership Annotations (Version v1) [Data set]. Zenodo. https://doi.org/10.5281/zenodo.18239580

Other public datasets used in this study, including the original Virtual KITTI 2 dataset, are available from their respective sources as cited in the manuscript.

For further inquiries, please contact the corresponding author.

## Conflict of Interest

None.

## Notes

https://doi.org/10.5281/zenodo.15375453

https://doi.org/10.5281/zenodo.16423201

https://doi.org/10.5281/zenodo.18239580

### Competing Interest Statement

The authors have declared no competing interest.

### Summary of Updates

Added public dataset availability information and made minor clarifications and organizational refinements to the text. The core theoretical framework, experimental results, and conclusions remain unchanged.

